# Strategic acyl carrier protein engineering enables functional type II polyketide synthase reconstitution *in vitro*

**DOI:** 10.1101/2023.08.02.551649

**Authors:** Kevin Li, Yae In Cho, Mai Anh Tran, Christoph Wiedemann, Shuaibing Zhang, Rebecca S. Koweek, Ngọc Khánh Hoàng, Grayson S. Hamrick, Margaret A. Bowen, Bashkim Kokona, Pierre Stallforth, Joris Beld, Ute A. Hellmich, Louise K. Charkoudian

## Abstract

Microbial polyketides represent a structurally diverse class of secondary metabolites with medicinally relevant properties and are synthesized by polyketide synthases (PKS). While Type I PKSs are large biosynthetic machineries composed of a single polypeptide chain, type II PKSs are minimally composed of a ketosynthase-chain length factor (KS-CLF) and a phosphopantetheinylated acyl carrier protein (*holo*-ACP) expressed separately. Although type II PKSs are found throughout the bacterial kingdom, and despite their importance to strategic bioengineering, type II PKSs have not been well-studied *in vitro*: In cases where the KS-CLF can be accessed via *E. coli* heterologous expression, the cognate ACPs are not activatable by the broad specificity *Bacillus subtilis* surfactin-producing phosphopantetheinyl transferase (PPTase) Sfp and, conversely, in systems where the ACP can be activated by Sfp, the corresponding KS-CLF is typically not readily obtained. Here, we report the high-yield heterologous expression of both cyanobacterial *Gloeocapsa* sp. PCC 7428 minimal type II PKS (gloPKS) components in *E. coli*, which allowed us to functionally reconstitute this minimal type II PKS *in vitro*. Initially, neither the cognate PPTase nor Sfp converted gloACP to its active *holo* state. However, by examining sequence differences between Sfp-compatible and -incompatible ACPs, we identified two conserved residues in gloACP that, when mutated, enabled high-yield phosphopantetheinylation of gloACP by Sfp. Using analogous mutations, other previously Sfp-incompatible type II PKS ACPs from different bacterial phyla were also rendered activatable by Sfp. This demonstrates the generalizability of our approach and breaks down a longstanding barrier to type II PKS studies and the exploration of complex biosynthetic pathways.

## Introduction

The promise of harnessing microbial biosynthetic machineries to gain sustainable access to structurally complex molecules with diverse medicinal properties has inspired decades of research.^1,2^ Polyketides represent a class of natural products with a particularly strong track record for benefiting human health. Landmark studies in the 1990s revealed that polyketide synthases (PKSs) are encoded by biosynthetic gene clusters (BGCs) and act as multifunctional protein assemblies responsible for the microbial biosynthesis of polyketides.^3,4^ In type I PKSs, all protein components required for substrate recognition, activation, transfer and release are organized in a single-chain multidomain protein complex.^5^ In contrast, the components of type II PKS are expressed separately as monofunctional entities, thereby presenting unique opportunities for strategic engineering.^6^ Aromatic polyketides produced by type II PKS are important targets for such efforts due to their widespread use as anticancer agents (*e*.*g*., doxorubicin) and antibiotics (*e*.*g*., tetracycline).^7^

In a minimal aromatic type II PKS, only two essential components are required: a ketosynthase-chain length factor (KS-CLF) and an acyl carrier protein (ACP), which serves to tether the growing polyketide chain and acetate building blocks. This minimal PKS catalyzes the decarboxylative Claisen-like condensations that convert acetate-based building blocks into a nascent β-ketoacyl intermediate.^7^ Once an ACP-bound β-ketoacyl intermediate reaches its programmed length (in part directed by the KS-CLF), it is transformed into the target natural product by a team of accessory enzymes.

The modularity of type II PKSs have made their *in vitro* functional reconstitution the focus of numerous research efforts.^6^ However, attempts to obtain intact KS-CLF enzymes through heterologous expression in fast-growing and easily accessible hosts such as *Escherichia coli* have been challenging, often because of lack of protein production or structural integrity.^8,9^ This has severely limited access to KS-CLFs, making it necessary to isolate them from slow-growing and often difficult to genetically manipulate actinomycete hosts.^10–22^ To address this problem, the focus recently shifted towards KS-CLFs from underexplored phyla inferred to be more closely related to the *E. coli* host than the actinomycete counterparts and thus presumed to be better suited for heterologous expression.^23–25^ Indeed, numerous cases of the successful heterologous expression of non-actinomycete type II PKS KS-CLFs in *E. coli* have been described, but always without obtaining the cognate ACP in its activated (“*holo*”) form.^26–29^ ACP activation occurs post-translationally with a phosphopantetheinyl transferase (PPTase) that covalently attaches a coenzyme A (CoA)-derived 4′-phosphopantetheine (Ppant) arm to a conserved serine residue on the *apo*-ACP.^30^ The *holo*-ACP’s Ppant arm then acts as a molecular tether to shuttle the building blocks and polyketide intermediates through the PKS during biosynthesis. In the laboratory, native or engineered^31^ surfactin-producing PPTase from *Bacillus subtilis* (Sfp) has typically been used to activate carrier proteins from diverse families including PKSs, fatty acid synthases (FASs) and non-ribosomal peptide synthetases.^30,32^ However, so far there has been limited success using Sfp to activate ACPs in high yields from non-actinomycete systems.^6,26,28^

Together, these two issues present a road block for the functional reconstitution of a minimal type II PKS, *i*.*e*., for systems where the ACP can be activated by Sfp, the KS-CLF is typically not readily obtained in high yields and where the KS-CLF can be accessed via *E. coli* heterologous expression, the cognate ACPs could not be activated by Sfp.^26,28^

Here, we used the cyanobacterial *Gloeocapsa* sp. PCC 7428 type II PKS (gloPKS) to overcome these difficulties. High yields for both the gloKS-CLF and the cognate *apo*-gloACP could be obtained from *E. coli* heterologous expression. However, similar to observations by others,^26,28^ gloACP could not readily be converted to the active *holo*-form by Sfp. By comparing related Sfp-compatible and -incompatible ACP sequences, we identified two residues important for Sfp activation. Mutagenesis of these residues in gloACP did not perturb ACP structure and enabled highly efficient phosphopantetheinylation by Sfp. This minimal mutagenesis strategy was conferrable to other Sfp-incompatible ACPs from other bacterial phyla, thus demonstrating the generalizability of the approach. In the case of the gloPKS, the *holo*-ACP could be loaded with a salicyl-priming unit and malonyl substrates. Together with the heterologously expressed and purified gloKS-CLF, this led to the functional reconstitution of this minimal type II PKS *in vitro* and the generation of a range of aromatic products. This study thus represents an important step towards the use of type II PKS for future bioengineering approaches.

## Results and Discussion

### *Gloeocapsa* sp. PCC 7428 minimal type II PKS components can be heterologously expressed in *E. coli* with high yields

Previous bioinformatic and phylogenetic analyses suggested that *Gloeocapsa* sp. PCC 7428 harbors a type II PKS BGC (Supplementary Figure S1) containing a *ks-clf* gene that is inferred to be more closely related to the *E. coli* FAS KS-encoding gene *fabF* than other actinobacterial *ks-clf*s.^23,24^ Together with the transcriptional coupling of the glo *ks* and *clf* genes in the native host, this indicated that gloKS-CLF might be a promising candidate for heterologous expression in *E. coli*.^26,28^ Indeed, we found that compared to the relatively slow-growing actinobacterial host, gloKS-CLF could be obtained in roughly 25-fold greater yields, *i*.*e*., with >20 mg/L purified protein from *E. coli* BL21(DE3) cells when expressed under a single promotor (Supplementary Figure. S2 and S3). Likewise, the cognate gloACP could be obtained in high amounts (∼17 mg/L) and was present exclusively in its *apo*-form upon purification (Supplementary Figure. S2 and S4).

### *Apo*-gloACP is not readily converted to its active *holo*-form

Typically, ACPs are obtained in their *holo*-form through either the heterologous expression in *E. coli* BAP1 cells, a genetically engineered derivative of the BL21 (DE3) strain carrying background expression of Sfp,^33^ or the *in vitro* reaction of purified ACP with recombinant *B. subtilis* Sfp.^31,34,35^ An Sfp with increased catalytic efficiency and substrate scope, termed Sfp R4-4, is typically used for ACP activation.^31^ We also made use of this variant and refer to it as Sfp throughout the manuscript. Neither expression in *E. coli* BAP1^33^ nor treatment of purified *apo*-ACP with Sfp yielded the phosphopantetheinylated gloACP, as shown by mass spectrometry (Supplementary Figures S4 and S5). Co-expression of gloACP with its cognate gloPKS BGC PPTase in *E. coli* also did not lead to the desired production of *holo*-ACP (Supplementary Figures S6 and 7). Finally, using an *in vitro* approach, we co-incubated *apo*-gloACP with purified gloPPTase, produced separately in *E. coli*, where it can be expressed at low levels (Supplementary Figure S8). Again, this did not yield *holo*-ACP (Supplementary Figure S9). Even in the presence of the putative CoA:ligase domain from *Gloeocapsa* (salicylate CoA:ligase; gloSCL; Supplementary Figures S1 and S2) and its cofactors (adenosine triphosphate (ATP) and salicylic acid), a strategy that resulted in ACP activation in other systems,^26,29^ the gloACP evaded activation (Supplementary Figure S10). These data highlight that gloACP activation is not straightforward and represents a barrier to the study of the minimal type II gloPKS.

### Sfp interacts with gloACP, but does not convert it to its *holo-*form

Initially, we wondered whether the inability of Sfp to activate gloACP might be due to structural differences between gloACP and ACPs known to be Sfp-compatible^*e*.*g*., 30,32^ For a structural characterization of gloACP, we determined the backbone and sidechain NMR assignments of the ^13^C, ^15^N-labeled protein (Supplemental Figure S11) and determined a chemical shift-derived structural model using CS-Rosetta^36^ (Figure 1A, Supplementary Figure S12). Comparison with available X-ray and NMR structures of Sfp-compatible ACPs from canonical type II FAS (*E. coli* AcpP, PDB: 1T8K), actinobacterial type II PKS (*Streptomyces coelicolor* ActACP, PDB: 2K0Y), and type I PKS (*Saccharopolyspora erythraea*, DEBS ACP2, PDB: 2JU2) systems revealed small structural differences but an overall canonical ACP fold for the *Gloeocapsa* protein (Figure 1B). GloACP differs from AcpP and ActACP by two short helical insertions between helix I and helix II, which we accordingly refer to as helix I′ and helix II′. Furthermore, gloACP lacks helix III’, an insertion between helix II and III seen in some other ACPs.^30^ Nonetheless, the overall secondary and tertiary structure arrangements of gloACP and importantly, the position of the serine residue at the *N*-terminus of helix II required for Ppant arm attachment is conserved compared to known Sfp substrates.

**Figure 1.**
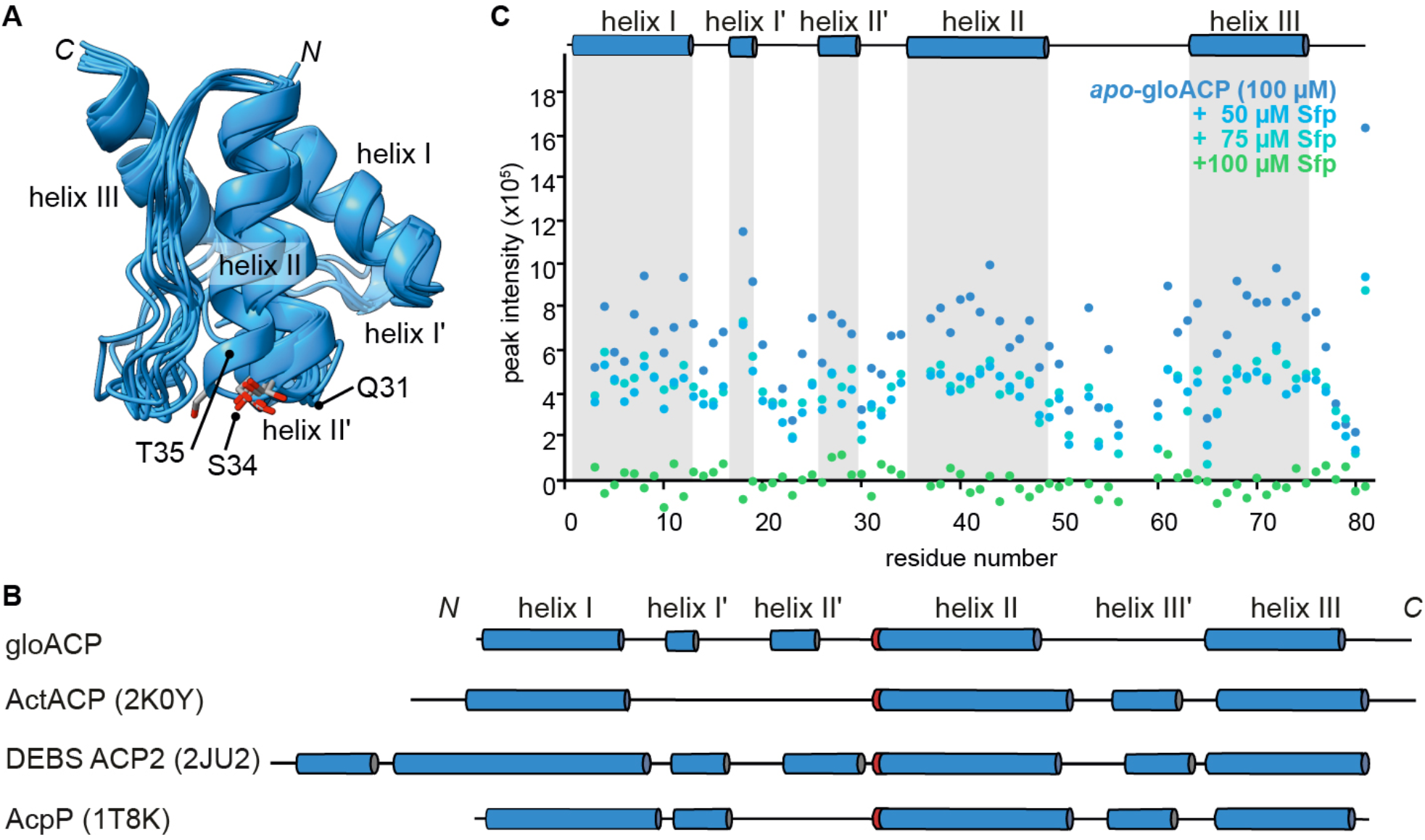
gloACP has a canonical fold and can interact with the *B. subtilis* PPTase Sfp. (A) The ten lowest energy structural models of WT gloACP obtained by CS-Rosetta. Residues Q31 and T35 important for Sfp-based ACP activation, as well as S34, the attachment point of the phosphopantetheine arm are marked. (B) Comparison of topology models for gloACP and available structures of Sfp-activatable, prototypical ACP domains: a type II PKS ACP (ActACP, PDB: 2K0Y) from *Streptomyces coelicolor*, a type I PKS ACP (DEBS ACP2, PDB: 2JU2) from *Saccharopolyspora erythraea*, and a type II FAS ACP (AcpP, PDB: 1T8K) from *Escherichia coli* (cylinders represent alpha-helices with red marking indicating the serine point of attachment of the Ppant arm. (C) NMR peak intensities of ^15^N-labeled gloACP upon Sfp titration. Severe line broadening at a 1:1 molar ratio (both proteins at 100 μM) indicates tight complex formation (see also Supplementary Figures S11–13).

Previously, we and others have successfully used NMR chemical shift perturbations to assess transient ACP-protein interactions.^37–39^ Here, we titrated the ^15^N-labeled 11 kDa gloACP with unlabeled 27 kDa Sfp and observed line broadening at sub-stoichiometric ratios, suggesting the formation of a larger molecular weight complex (Figure 1C, Supplementary Figure S13). At an equimolar ratio, all gloACP signals disappeared, consistent with tight complex formation between gloACP and Sfp. Together this indicates that neither structural differences nor lack of protein interactions are likely to be the decisive cause for the inability of Sfp to activate gloACP.

### Minimal gloACP sequence modifications enable Sfp-mediated transformation from *apo* to *holo-*form

To overcome the inability of Sfp to activate the carrier protein, we looked at the tyrocidine A synthetase peptidyl carrier protein (TycC3_PCP) from *Bacillus brevis*, which has been studied in complex with Sfp.^40^ TycC3_PCP residues G42, L46, and M49 were found to be crucial for phosphopantetheine transfer by Sfp. A multiple sequence alignment of Sfp-compatible and incompatible carrier proteins suggests that two out of the three positions identified in TycC3_PCP may also affect Sfp (in)compatibility with gloACP (Figure 2A, Supplementary Figure S14). In Sfp-compatible ACP sequences, the position of the glycine and a hydrophobic residue analogous to TycC3_PCP L46 are generally preserved (marked in cyan in Fig 2A). In contrast, these residues are more varied in Sfp incompatible ACPs with *e*.*g*., gloACP Q31 and T35 (marked in magenta in Figure 2A). These residues are within helix II and structurally in close proximity to the serine important for phosphopantetheinylation (Figure 1A). To evaluate whether these residues play a role in Sfp-based activation, we generated single and double gloACP mutants emulating the TycC3_PCP sequence, *i*.*e*., gloACP^Q31G^, gloACP^T35L^ and gloACP^Q31G/T35L^. All gloACP mutants could be heterologously expressed in *E. coli* and purified in their *apo*-form (Supplementary Figures S15 and S17). As described above, WT gloACP was not phosphopantetheinylated by Sfp *in vitro* (Supplementary Figure S5). Likewise, no phosphopantetheinylation for gloACP^Q31G^ and minimal conversion of gloACP^T35L^ (∼ 5% conversion after 16h) by Sfp were observed by mass spectrometry (Supplementary Figure S18). In contrast, gloACP^Q31G/T35L^ was successfully converted to its *holo*-form, yielding ∼95% conversion (Supplementary Figure S16). Interestingly, this gain in Sfp-compatibility for gloACP^Q31G/T35L^ was not due to structural changes compared to the WT ACP, as indicated by the minimal chemical shift differences of the two proteins, but rather due to an apparent loosening of the interaction with Sfp as indicated by the lower degree of line broadening (Figure 2B, C).

**Figure 2.**
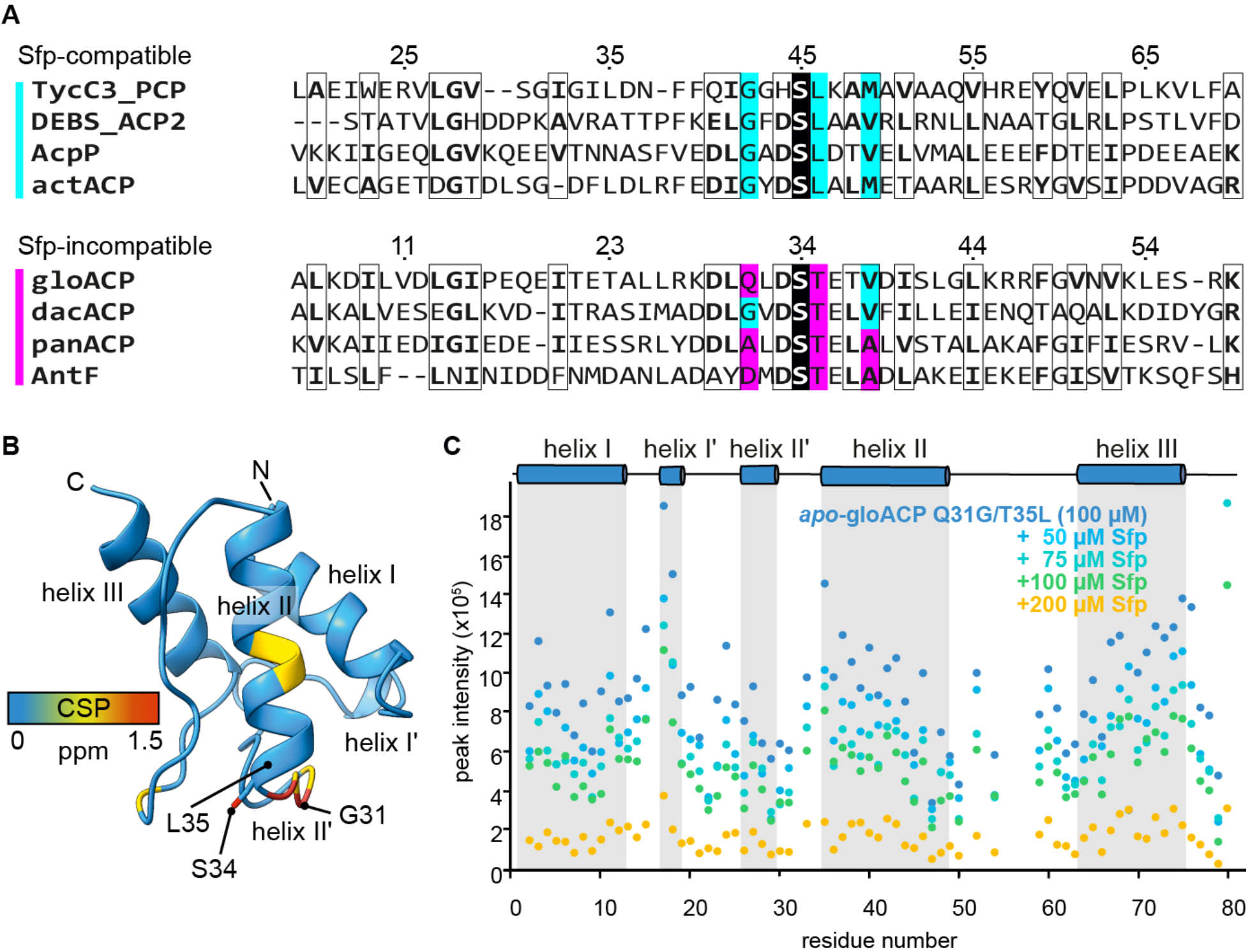
Sfp compatibility can be inferred from ACP sequence, enabling targeted modifications to render ACPs activatable through loosened Sfp interactions. (A) Multiple sequence alignment of carrier proteins known to be modified by Sfp (top four with cyan bar) and non-actinobacterial Sfp-incompatible ACPs (bottom four with magenta bar). Numbering is based on the TycC3_PCP^40^ and gloACP sequences, respectively. GloACP, dacACP, and panACP were used in this study (see main text for details), while AntF is a previously characterized non-actinobacterial ACP from *Photorhabdus luminescens*.^41^ The serine carrying the phosphopantetheine arm is highlighted in black in each sequence. Black boxes represent residues containing greater than 70% physicochemical similarity. Based on our and previously published functional data (see main text for details) as well as the sequence alignments, we hypothesized that the highlighted residues (cyan and magenta, respectively) direct Sfp-(in)compatibility. (B) Chemical shift differences between wild-type (WT) gloACP and gloACP^Q31G/T35L^ (Supplementary Figure S11 and S12) mapped on the CS-Rosetta model of WT gloACP show minimal changes upon mutation. (C) Relatively higher peak intensity of ^15^N-labeled gloACP^Q31G/T35^ (100 μM) upon titration with unlabeled Sfp indicates looser interaction compared to WT gloACP (see Figure 1C, Supplementary Figure S13).

To reflect the presence of other hydrophobic residues in some of the available ACP sequences at the position homologous to gloACP T35, we also generated the Q31G/T35I double mutant. Sfp successfully phosphopantetheinylated this variant, albeit with lower efficiency than the Q31G/T35L double mutant (∼30% conversion, Supplementary Figure S19). Importantly, minimal sequence modifications convert an Sfp-incompatible into a compatible ACP.

### Minimal sequence adaptations present a general route to render previously inaccessible ACPs Sfp-compatible

Having identified the two relevant residues for the cyanobacterial gloACP activation by Sfp, we next evaluated the generalizability of this approach for other ACPs. To this end, we selected type II PKS ACPs from *Delftia acidovorans* (dacACP) and *Paenibacillus* sp. FSL R7-277 (panACP) (Figure 2A), representing the proteobacteria and firmicutes phyla.

Both dacACP and panACP contain a threonine residue at positions analogous to gloACP T35 and TycC3_PCP L46 (Fig 2A). According to our hypothesis, these residues should interfere with Sfp-compatibility. In addition, panACP contains two alanine residues (A31, A38) at positions analogous to TycC3_PCP G42 and M49, that we would thus deem to hamper Sfp-compatibility. In contrast, we would predict the respective residues dacACP, *i*.*e*., G31, V38, to enhance Sfp-compatibility, suggesting that Sfp may retain some ability to modify dacACP.^40^ Both WT dacACP and panACP were purified from *E. coli* in their *apo*-form (Supplementary Figures S20 and S22).

In agreement with the expectation for a tapered ability of either ACP to become activated by Sfp derived from the sequence comparison, we saw that WT panACP was not phosphopantetheinylated, while WT dacACP was phosphopantetheinylated to a very low degree (< 5%, Supplementary Figures S20–23). Next, we aimed to transform panACP and dacACP into fully Sfp-compatible proteins. Analogous to the gloACP^Q31G/T35L^ variant, we generated two constructs emulating the TycC3_PCP sequence expected to enable phosphopantetheinylation, *i*.*e*., dacACP^T43L^ and panACP^A30G/T34L/A37V^. Both variants were successfully purified from *E. coli* in their *apo*-form (Supplementary Figures S24 and S26) and found to be fully phosphopantetheinylated by Sfp (Supplementary Figures S25 and S27). This underscores the importance of a nonpolar residue in a position analogous to TycC3_PCP L46 for efficient phosphopantetheinylation by Sfp. Importantly, these findings also suggest that Sfp (in)compatibility can be directly predicted from the ACP sequence and minimal protein engineering can endow type II PKS ACPs with the ability to be modified by Sfp. Together with the increased access to KS-CLFs via *E. coli* heterologous expression, this opens the door for the functional reconstitution of hitherto inaccessible type II PKSs *in vitro*.

### Functional reconstitution of a minimal type II PKS *in vitro*

After gaining access to the quantitative modification of gloACP by Sfp, we aimed to functionally reconstitute the minimal type II gloPKS composed of ACP and KS-CLF *in vitro* (Figure 3). We first assessed whether the Ppant arm of *holo*-gloACP^Q31G/T35L^ can be acylated with a malonyl group, a critical step in the formation of a growing polyketide chain. The majority of ACPs studied to date require malonyl-CoA:acyl carrier protein transacylases (MCATs) to catalyze this process.^42,43^ However, in select cases, MCATs are not required and can ‘self-malonylate’.^13,44^ Upon incubation with malonyl CoA, a low degree of self-malonylating activity was observed for *holo*-gloACP^Q31G/T35L^ (Supplementary Figure S28). However, for full conversion from *holo-* to malonyl-gloACP^Q31G/T35L^, co-incubation with MCATs from either *Streptomyces coelicolor* (ScFabD) or *E. coli* (EcFabD) was required (Supplementary Figures S29 and S30).

**Figure 3.**
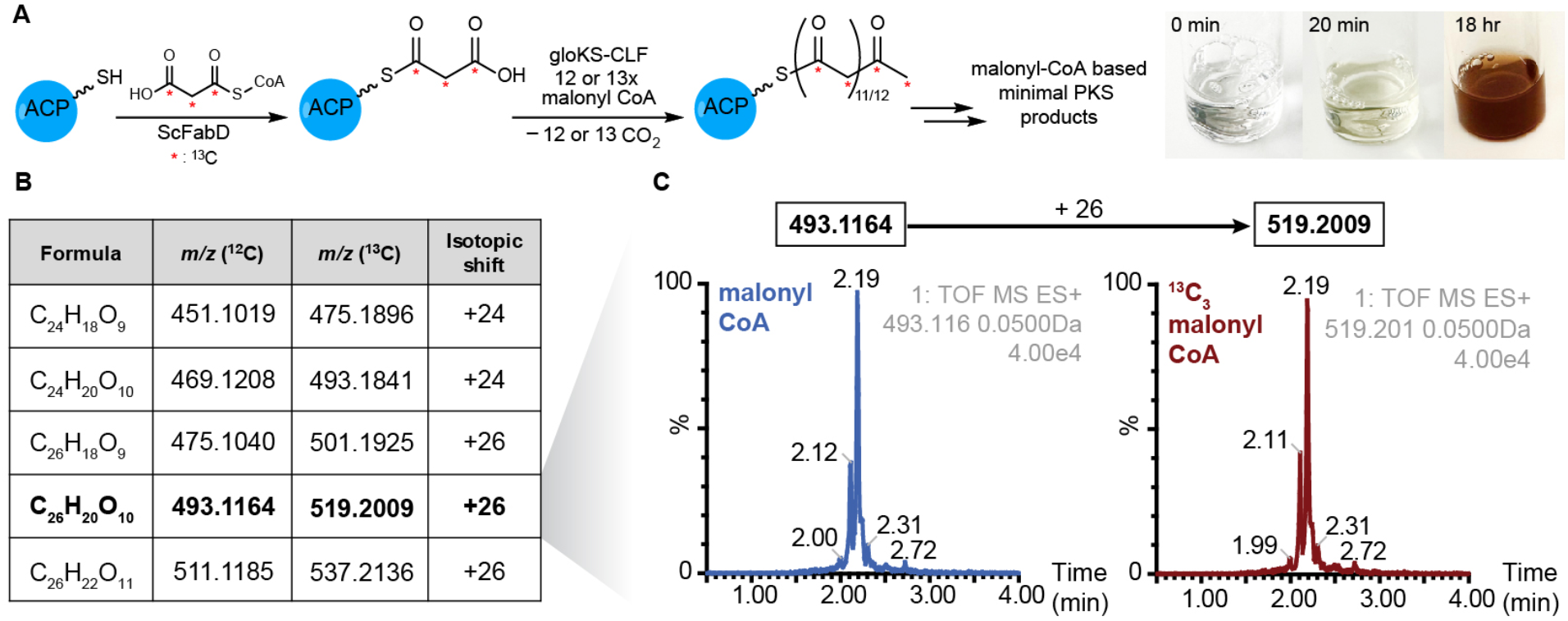
Reconstitution of holo-gloACP^Q31G/T35L^ and gloKS-CLF function yield a functional minimal type II PKS *in vitro*. (A) Proposed *in vitro* reaction scheme of gloPKS polyketide production by the minimal gloPKS components and color switch of reaction mixture after addition of ScFabD and malonyl CoA to *holo*-gloACP and gloKS-CLF. (B) Chemical formulas and experimental m/z values (^12^C and ^13^C) of putative precursor polyketide metabolites produced by the minimal gloPKS with malonyl CoA. (C) Extracted ion chromatograms of the putative metabolite C_26_H_20_O_10_. ^13^C_3_-malonyl-CoA supplementation leads to a mass shift of +26 amu in both cases (For additional extracted ion chromatograms of putative precursor polyketides see Supplementary Figure S31).

Next, we explored whether the minimal gloPKS can produce polyketides *in vitro*. This requires the productive interaction of both heterologously produced protein units to mediate the two central polyketide biosynthesis reactions: chain transfer and elongation. Consistent with the formation of polyaromatic products, mixing gloKS-CLF, *holo*-gloACP^Q31G/T35L^, ScFabD and malonyl CoA resulted in in the generation of compounds and a concomitant color change from clear to bright orange/yellow (Figure 3A, right). Isolation of the colored molecules and analysis by high-resolution mass spectrometry showed the presence of compounds with molecular formulas of C_24_H_18_O_9_ (*m/z* = 451), C_24_H_20_O_10_ (*m/z* = 469), C_26_H_18_O_9_ (*m/z* = 475), C_26_H_20_O_10_ (*m/z* = 493), and C_26_H_22_O_11_ (*m/z* = 511) (Fig 3B and Supplementary Figure S31). Isotopic shift experiments feeding with ^13^C_3_-malonyl-CoA resulted in isotopic shifts of 24-amu (451, 469) and 26-amu (475, 493, 511), suggesting the production of dodecaketides and tridecaketides via condensation of 12 and 13 molecules of malonyl-CoA, respectively by the minimal gloPKS *in vitro* (Figure 3C). The successful activation of the PKS biosynthetic machinery led to the production of a highly complex product mixture, precluding the purification of individual compounds in sufficient quantities for additional structural characterization.

### Additional enzymes and substrates expand the scope of the reconstituted minimal gloPKS

After obtaining a functional minimal gloPKS *in vitro*, we wondered whether its product scope can be expanded through addition of alternative substrates. The gloPKS BGC (Supplementary Figure S1) shows similarities to the thermorubin-producing PKS from *Laceyella sacchari*, suggesting that the gloPKS can be primed by salicylic acid (Supplementary Table S1).^46^ Indeed, we observed that the addition of ATP and the heterologously expressed cognate salicylate:CoA ligase (gloSCL) allowed priming of the *holo*-gloACP^Q31G/T35L^ with salicylic acid (Supplementary Figure S32). In the presence of the gloKS-CLF, salicyl-gloACP^Q31G/T35L^ underwent multiple rounds of chain extension with the malonyl-CoA building block, resulting in C29 salicyl-primed products (Figure 4A and 4B). High-resolution mass spectrometry revealed the formation of products consistent with the formulas C_29_H_20_O_10_ and C_29_H_22_O_11_, *i*.*e*., *m/z* 529 and 547 [M+H]^+^.

**Figure 4.**
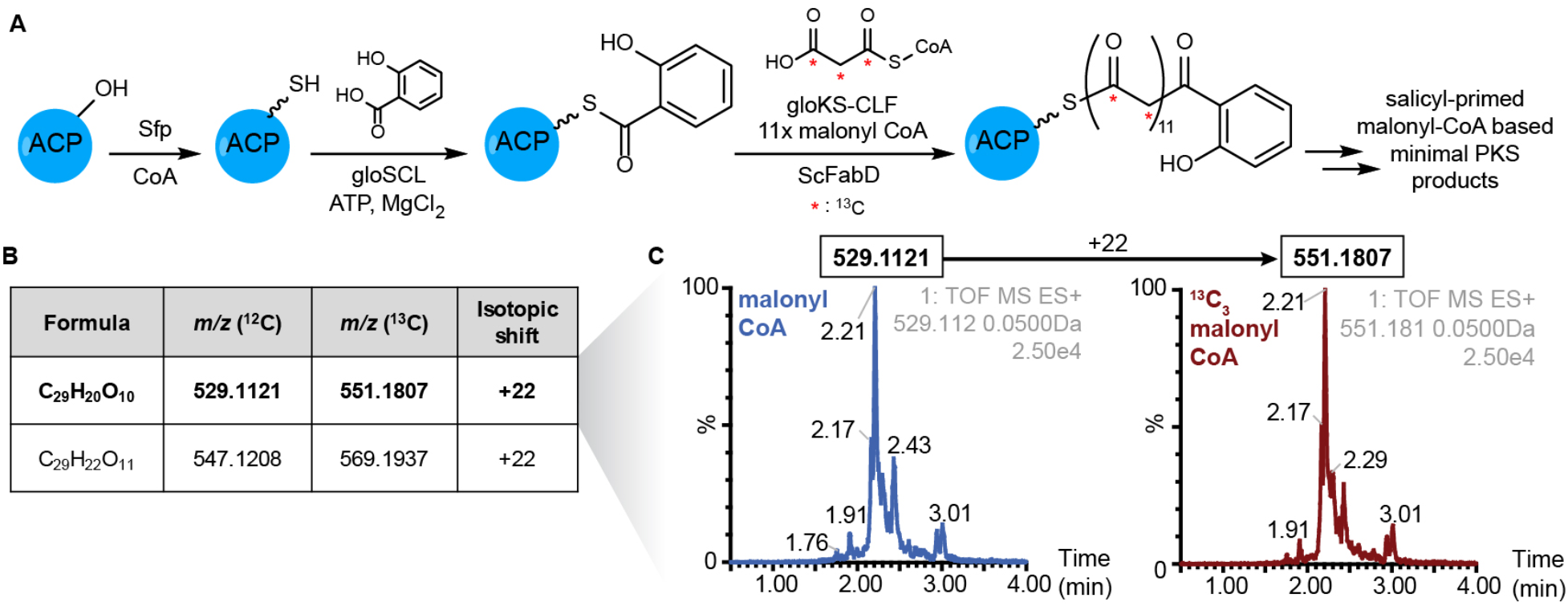
Salicyl priming expands the product profile of the reconstituted minimal gloPKS. (A) Proposed *in vitro* gloPKS polyketide production scheme. Red stars represent carbon atoms derived from malonyl-CoA and anticipated to be labeled during ^13^C_3_-malonyl-CoA isotope experiments. (B) Chemical formulas and experimental *m/z* values (^12^C and ^13^C) of putative precursor polyketide metabolites obtained from LC-MS experiments. (C) Extracted ion chromatograms of the putative metabolite C_29_H_20_O_10_. Upon ^13^C_3_-malonyl-CoA supplementation, a mass shift of +22 amu is observed (for additional extracted ion chromatograms of putative precursor polyketide products see Supplementary Figure S33).

Feeding of ^13^C_3_-labeled malonyl-CoA led to a mass shift of 22 amu for both products (Figure 4B, C, Supplementary Figure S33), indicating the incorporation of 11 malonyl-CoA units. This is consistent with the thermorubin biosynthetic pathway, which affords a 29-carbon polyketide backbone from seven carbons derived from salicylate and 22 carbons derived from malonyl-CoA.^45,46^

Importantly, these results show that the engineered *holo*-gloACP^Q31G/T35L^ variant remains a functional unit within an expanded PKS roster that includes pathway priming components. Moreover, the distinct products manufactured in the presence versus absence of salicylic acid priming supports the concept that core type II PKS biosynthetic components can be shuffled to direct the chain lengths of polyketides produced.

## Conclusions

Access to components of minimal type II PKS opens the door for investigations focused on substrate specificity and screening, kinetics, mechanisms, and structural analyses but was historically hindered by the lack of access to inactive ACP in conjugation with the cognate KS-CLF. Here, we have shown that ACP compatibility with the PPTase Sfp, and by extension ACP activation for the functional reconstitution of type II PKSs, can be directly inferred from the ACP sequence. Minimal protein engineering of the ACP enables the efficient transformation of heterologously produced non-actinomycete *apo*-ACPs into their active *holo*-form by Sfp. This, together with improved access to KS-CLFs via the *E. coli* heterologous expression of non-actinomycete target genes as outlined in the recent literature^26–29^ and highlighted in the current work, allowed to reconstitute a functional minimal type II PKS *in vitro*. Using the minimal gloPKS, we saw that this system can make use of diverse priming molecules to create a large set of aromatic products. However, it remains unclear whether the product diversity observed *in vitro* aligns with the *in vivo* output profile. It is tempting to speculate that unknown crowding factors or accessory proteins may regulate product synthesis specificity. Alternatively, the ability to incorporate varying priming molecules and the resulting product promiscuity may also be a feature of synthases in bacteria, as these organisms are exposed to fast changing environmental demands (*e*.*g*., nutrient fluctuations or encounters with competition). The *Gloeocapsa* type II PKS is particularly well-suited to follow up on important questions that require bridging *in vitro* and *in vivo* aspects. The organism is culturable under laboratory conditions^47^ and, as we have shown in the present work, the gloPKS can be heterologously produced and functionally reconstituted *in vitro*. Future work will focus on understanding the molecular details of the interplay between transient domain interactions and biosynthetic steps.

The workflow presented here provides a general, accessible, and easy route to *holo*-ACP from biosynthetic machineries in diverse phyla that were hitherto inaccessible. This progress will enable future exploits into combinatorial biosynthesis schemes with reconstituted type II PKS domains. It is intriguing to point out that an apparent loosening enabled efficient phosphopantetheinylation of the ACP by Sfp. This suggests that the lack of activation of the WT gloACP to *holo-*form is due to a lack of catalysis rather than lack of binding. These findings showcase another example for the importance of transient, short-lived interactions in functional PKS modules.^21,39,48–50^ Enhancing intramolecular contacts that, as we have seen, can be detrimental to function, could thus guide the development of inhibitors targeting bacterial PPTases implicated in common pathogens’ associated virulence factors, further expanding the impact of type II PKS research.^32,51^

## Methods

### Native Organism Culturing and DNA Isolation

*Gloeocapsa* sp. PCC 7428 (ATCC^®^ 29159™) was purchased as a liquid culture from the American Type Culture Collection (ATCC^®^) and inoculated in 30 mL of BG-11 liquid medium (ATCC^®^ medium 616).^47^ Cultures were grown at room temperature with indirect light exposure for 6–7 weeks with aeration and no shaking. Genomic DNA was isolated following the protocol described in Saha *et al*.^52^

### Molecular cloning

*E. coli* DH5α competent cells were used for all cloning experiments. AntiSMASH 5.0 was used to identify genes corresponding to *Gloeocapsa* sp. PCC 7428 (ID: CP003646) ACP, KS-CLF, PPTase, and SCL genes.^53^ Each gene was amplified from purified genomic DNA using primers listed in Supplementary Table S2 and standard Q5 High-fidelity DNA polymerase. For genes corresponding to ACPs from *Paenibacillus* sp. FSL R7-277 and *Delftia acidovorans*, codon optimized DNA was purchased from Twist Bioscience and amplified (see Table S2 for DNA sequences)Gel-purified amplimers were inserted into linearized pET28a vector backbone between NdeI and EcoRI cleavage sites (except for the malonyl-CoA synthetase, MatB, which was inserted between NdeI and XhoI cleavage sites) via Gibson assembly^54^ using an NEB Q5 high-fidelity 2X master mix kit (for plasmids, see Supplementary Table S2). The expression plasmid for EcFabD was provided as a gift from the Campopiano Research group at University of Edinburgh, the plasmid for ScFabD was purchased from Gene Universal. Plasmid sequences were confirmed by whole plasmid sequencing (Plasmidsaurus).

### Multiple sequence alignment (MSA)

Amino acid sequences of the carrier proteins of interest were aligned using Clustal Omega and ESPript 3.0.^55,56^

### Site directed mutagenesis

Mutant plasmids of the ACPs were obtained using the NEB Q5 site directed mutagenesis kit following the manufacturer’s protocol. In brief, amplification was performed on the mixture of Q5 Hot Start High-Fidelity 2X Master Mix, 10 μM forward primer, 10 μM reverse primer, template DNA (1-25 ng of the WT ACP plasmid), and nuclease-free water. Kinase, ligase & DpnI (KLD) treatment was conducted by mixing the PCR product, 2X KLD reaction buffer, 10X KLD enzyme mix, and nuclease-free water, followed by incubation at room temperature for 5 minutes. The KLD mix was transformed into *E. coli* DH5α competent cells and plated onto LB agar supplemented with 50 μg/mL kanamycin (Kan). pGloACP was amplified with four distinct primer sets to create the gloACP mutants gloACP^Q31G/T35L^, gloACP^Q31G/T35I^, gloACP^Q31G^, and gloACP^T35L^. Vectors pPanACP and pDacACP encoding WT panACP and dacACP, respectively, were used to create mutants panACP^A30G/T34L/A37V^ and dacACP^T43L^. All primers and insert sequences are provided in Supplementary Table S2.

### Protein expression & purification

All proteins were purified according to previously reported protocols.^57^ In brief, plasmids were transformed into competent *E. coli* BL21(DE3) cells or, to aim to express *holo*-gloACP, into BAP1 cells^31^ as specified in the main text. Seed cultures were prepared by inoculating a single bacterial colony in 10 mL of LB supplemented with 50 μg/mL Kan and incubating overnight at 37 °C with shaking for 18 hours. Production cultures (1 L LB supplemented with 50 μg/mL Kan) were inoculated with the 10-mL overnight seed culture, incubated at 37 °C and induced with isopropyl β-D-1-thiogalactopyranoside (IPTG, final concentration of 250 μM) once cultures reached an OD_600_ of 0.4–0.6. After overnight incubation (18 °C, 200 rpm gentle shaking), cells were harvested by centrifugation (4424 × *g*, 4 °C, 20 minutes) and stored at –80 °C until purification. For purification, cell pellets were resuspended on ice using lysis buffer (50 mM sodium phosphate buffer, 10 mM imidazole, 300 mM NaCl, 10% (v/v) glycerol, pH 7.6) and lysed (Microson XL-200 Ultrasonic Processor: 10 × 30 sec pulses with 30 sec rest in between at 40% amplitude). Cellular debris was removed via centrifugation (17,000 × *g*, 4 °C, 1 hour), and the resulting supernatant was transferred to nickel-NTA agarose equilibrated with lysis buffer and mixed for 1–2 hours at 4 °C with gentle rocking. The protein-bound resin suspension was transferred into a gravity flow column (Bio-Rad Econo-Pac^®^-chromatography column) equipped with a 30 μM polyethylene bed and allowed to settle before collecting the flow through and washing twice using wash buffer (2 × 100 mL of 50 mM sodium phosphate buffer, 30 mM imidazole, 300 mM NaCl, 10% (v/v) glycerol, pH 7.6). Finally, the His_6_-tagged proteins were eluted with 10 mL of elution buffer (50 mM sodium phosphate buffer, 250 mM imidazole, 100 mM NaCl, 10% (v/v) glycerol pH 7.6) and quantified using UV/VIS or bicinchoninic acid (BCA) assays for proteins lacking amino acids with aromatic side chains. All proteins were aliquoted, flash-frozen, and stored at –80 °C. Sodium dodecyl sulfate polyacrylamide gel electrophoresis (SDS-PAGE) and Western Blot were used to confirm protein purity (Detailed in Supplementary Methods, Figures S2 and S8).

### NMR spectroscopy

Uniformly ^15^N-labeled and ^13^C/^15^N labeled *N*-terminal His_6_-tagged gloACP was expressed in *E. coli* BL21(DE3) gold cells grown. The cells were grown in M9 minimal medium^58^ supplemented with 1 g/L ^15^NH_4_Cl and 2 g/L U-^13^C-glucose as the sole nitrogen and carbon sources, respectively, following a spin-down protocol. For ^15^N labeling, 1 g/L ^15^NH_4_Cl was used as the sole nitrogen source and the M9 minimal medium was supplemented with unlabeled glucose (2 g/L) as carbon source. Transformed *E. coli* cells were first grown in 1 L LB medium at 37 °C with shaking at 180 rpm to an OD_600_ = 0.4–0.6 and then centrifuged (25 °C, 5000 x *g*, 10 min). The cell pellets were resuspended in 1 L M9 minimal medium supplemented with 50 μg/mL Kanamycin and incubated for 30 min at 37°C with shaking at 180 rpm. Protein expression was induced with 312 μM IPTG and cells were grown for 16 hours at 18 °C with shaking at 180 rpm. Cells were then harvested via centrifugation (4 °C, 5000 x g, 20 min).

Cell pellets were resuspended in 100 mL of 50 mM sodium phosphate buffer (pH 7.6) containing 300 mM NaCl, 10 mM imidazole, 10% (v/v) glycerol, a protease inhibitor mix (1:1000 dilution), and a spatula tip of lysozyme, DNase, and RNase. Cells were lysed by sonication, and the lysate was clarified by centrifugation (10,000 x g, 30 min, 4 °C). The supernatant was incubated with pre-equilibrated nickel-NTA beads (2 mL/L of culture) for 1 hour on an end-over-end rotator. The mixture was transferred to a gravity column, washed sequentially with 10 column volumes (CV) of 50 mM sodium phosphate buffer (pH 7.6) containing 100 mM NaCl, 10 mM imidazole, and 10% (v/v) glycerol, followed by 10 CV of the same buffer containing 30 mM imidazole. The protein was eluted in five 1 CV steps with 50 mM sodium phosphate buffer (pH 7.6) containing 100 mM NaCl, 250 mM imidazole, and 10% (v/v) glycerol. Protein-containing fractions were identified using a BCA assay, pooled, and concentrated using Sartorius Vivaspin 3000 MWCO filters. Size exclusion chromatography was performed using a HiLoad prep grade 16/60 Superdex 75 column equilibrated with 50 mM sodium phosphate buffer (pH 7.6) containing 100 mM NaCl.

All NMR experiments were conducted at 298 K on Bruker AVANCE III HD 600 MHz spectrometers equipped with cryogenic triple resonance probes (Bruker BioSpin GmbH, Rheinstetten). Triple resonance assignment experiments (HNCO, HN(CA)CO, HNCA, and HNCACB) were recorded using standard Bruker TopSpin pulse sequences for resonance chemical shift assignments. Both wild-type gloACP and gloACP^Q31G/T35L^ were measured at concentrations of 300–410 μM in 50 mM sodium phosphate buffer (pH 7.5) containing 100 mM NaCl, 5% (v/v) d_6_-glycerol. Samples were supplemented with 10% D_2_O for field-frequency locking and 0.1 mM 2,2-dimethyl-2-silapentane-5-sulfonic acid (DSS) for chemical shift referencing. The ^1^H signal of DSS was set to 0 ppm for direct referencing and the ^13^C and ^15^N chemical shifts were indirectly referenced according to their magnetogyric ratios.^59^ Data processing was performed using Bruker TopSpin 3.2 or 4.1 software, and analysis of wild-type gloACP, gloACP^Q31G/T35L^, and *holo*-gloACP^Q31G/T35L^ was carried out using CCPN V3.1.^60^ For titration experiments with Sfp R4-4, ^15^N-labeled WT gloACP and ^15^N-labeled gloACP^Q31G/T35L^ were used. NMR samples were prepared by diluting the stock solutions of ^15^N wild-type gloACP or ^15^N gloACP^Q31G/T35L^ (both 270 μM) in the appropriate buffer to achieve a final concentration of 100 μM gloACP or gloACP^Q31G/T35L. 1^H-^15^N correlation spectra were recorded for both proteins. For titration with Sfp R4-4, stocks of unlabeled Sfp (830 μM) were mixed with ^15^N-labeled ACP, 10% D_2_O and the appropriate amount of buffer. Dilution effects never exceeded 10% and were accounted for during data analysis.

### CS-Rosetta modeling

Three-dimensional gloACP structural models were calculated with CS-Rosetta^36^ (version 2.01rev2019.06 using 40 CPUs within the NMRbox virtual environment)^61^ using experimental backbone ^13^CO, ^13^C^α, 13^C^β, 1^H^α, 1^H^N^ and ^15^N^H^ chemical shift restraints. Before setting up the CS-Rosetta run, the referenced gloACP backbone chemical shifts were submitted to TALOS-N^62^ for generation of dihedral restraints and predicting highly flexible or disordered regions. Since TALOS-N predicted highly dynamic amino acid stretches at the *N*- and *C*-termini of the gloACP construct the *N*-terminal purification tag (residue -19–0) and the last five C-terminal residues (76–80) were removed for the modeling procedure. To facilitate proper folding, repulsive energy was used for those residues during structure calculation. GloACP residues 54–58 were included in the calculation run although classified as dynamic by TALOS-N.

The CS-Rosetta run was considered to have converged based on the backbone C^α^-atom RMSD (root mean square deviation) < 2 Å between the lowest re-scored energy structures. 1292 out of the 10,000 calculated structural models have a RMSD < 2 Å to the lowest energy model (Figure S12). The 10 lowest re-scored energy models with a mean C^α^-RMSD of 0.84 ± 0.33 Å were chosen.

### *In vitro* ACP phosphopantetheinylation assay

Standard phosphopantetheinylation assays of *apo*-ACPs (gloACP, panACP variants and dacACP variants) by Sfp were carried as previously described.^35,57^ In brief, 500 μL reactions were set up in 50 mM sodium phosphate buffer, pH 7.6 in glass vials at room temperature overnight with final concentrations of 150 μM ACP, 1 μM Sfp R4-4, 2.5 mM DTT, 1.5 mM CoA (lithium salt from CoALA Biosciences), and 10 mM MgCl_2_. Phosphopantetheinylation assays with *apo*-gloACP and gloPPT were set up under the same conditions, but with a gloPPT concentration of ∼2.5–10 μM and WT gloACP concentration of ∼200 μM. GloPPT phosphopantetheinylation reactions involving the addition of gloSCL and co-factors contained additional final concentrations of 3 μM gloSCL, 5 mM ATP (disodium trihydrate, Gold Biotechnology), and 0.5 mM sodium salicylate. The degree of conversion from *apo*-ACP into *holo*-ACP was assessed by LC-MS as described below.

### Analysis of ACPs by LC-MS

The loaded state of ACPs (*apo, holo, malonyl, salicyl*) of was analyzed by LC-MS (Supplementary Table S3 and Figures S4–7, 9, 10, 15–30, 32).^63,64^ For LC-MS analysis, protein samples were diluted to ∼10 μM using HPLC grade water and analyzed using an Agilent Technologies InfinityLab G6125B LC/MS coupled with an Agilent 1260 Infinity II LC system loaded with a Waters XBridge Protein BEH C4 reverse phase column (300 Å, 3.55 μm, 2.1 mm x 50 mm) heated to 45 °C. Samples were analyzed via electrospray ionization mass spectrometry (ESI-MS) in the positive mode. The following gradient was used: 0–1 min 5% B; 1–3.1 min 95% B; 3.1–4.52 min 95% B; 4.52–4.92 min 5% B; 4.92–9 min 5% B (solvent A = water + 0.1% (v/v) formic acid; solvent B = acetonitrile + 0.1% (v/v) formic acid). ACPs (regardless of loaded state) eluted with a retention time of 4.2-4.6 min, as observed by absorbance at 254 and 280 nm as well as total ion count). Following data collection, mass spectra were deconvoluted using ESIprot online^65^ and plotted using Origin (Version 8.60. OriginLab Corporation, Northampton, MA). The observed deconvoluted MS was compared to the calculated *M*_*W*_ for ACPs in their various loaded states. The relative intensity of the *m/z* peaks assigned to *apo* versus *holo* state was used to approximate the percent conversion.

### Tandem proteolysis mass spectrometry

Tandem proteolysis was performed on the SDS PAGE bands corresponding to the putative KS and CLF to confirm the protein identities by the Proteomics and Metabolomics Facility of the Wistar Institute located in Philadelphia, PA. Samples were reduced with tris (2-carboxyethyl) phosphine (TCEP), alkylated with iodoacetamide, and digested with trypsin. Resulting tryptic peptides were injected into a Waters ACQUITY UPLC Symmetry C18 trap column (100 Å, 180 μm x 2 mm packed with 5 μm C18 particles) and separated using their standard 90-minute gradient formed by solvent A (water + 0.1% (v/v) formic acid) and solvent B (acetonitrile + 0.1% (v/v) formic acid). Peptides were scanned from 400 to 2000 *m/z* under positive ionization. The full scan was conducted at 70,000 resolution and data-dependent MS/MS scans at 175,000 resolution were performed on the resulting ions that exceeded a minimum threshold of 20,000. Peptide match was set as “preferred, exclude isotopes” and charge-state screening was enabled to reject singly and unassigned charged ions. The resulting peptide sequences were identified using MaxQuant 1.6.1.0 software (Supplemental Figure S3).

### Malonylation of *holo*-gloACP^Q31G/T35L^

Malonylation protocols were adapted from Beltran-Alvarez *et al*.^66^ All reaction components were stored in 50 mM sodium phosphate buffer, pH 7.6. Prior to the addition of malonyl-CoA, *holo*-gloACP^Q31G/T35L^ and TCEP-HCl were mixed in a clear 2-mL glass vial and pre-incubated at 30 °C for 30 minutes without shaking. Final reaction concentrations were as follows: 75 μM *holo*-ACP, 1 mM TCEP-HCl, and 1 mM malonyl-CoA (lithium salt from CoALA Biosciences) in a total volume of 100 μL in 50 mM phosphate buffer, pH 7.6. For reactions involving EcFabD or ScFabD, these MCATs were added to reach final concentrations of 1.5 μM. Once all components were combined, reactions were incubated at 30 °C for 30 minutes prior to analysis of the ACP loaded state by LC-MS, as described above.

### Salicyl priming of *holo*-gloACP^Q31G/T35L^

Salicyl priming of *holo*-gloACP^Q31G/T35L^ was carried out in 50 mM sodium phosphate buffer, pH 7.6. The protein was pre-incubated with 1 mM TCEP-HCl for 30 minutes at 30 °C in a clear 2-mL glass vial before adding additional components. Final reaction concentrations were as follows: 75 μM *holo*-gloACP^Q31G/T35L^ 1 mM TCEP-HCl, 3 μM gloSCL, 0.5 mM sodium salicylate, 5 mM MgCl_2_, and 5 mM ATP in a total volume of 200 μL. Once all components were combined, reactions were incubated at 30 °C for 30 minutes prior to analysis of the ACP loaded state by LC-MS, as described above.

### gloPKS *in vitro* reactions with acetate priming

*In vitro* reaction protocols were adapted from Cheng *et al*.^21^ All reaction components were stored in 50 mM sodium phosphate buffer (pH 7.6) with 10% v/v glycerol. Prior to the addition of gloKS-CLF, ScFabD and ^12/13^C malonyl-CoA, *holo*-gloACP^Q31G/T35L^ and TCEP-HCl were mixed in a clear 2-mL glass vial and pre-incubated at 30 °C for 30 minutes without shaking. Final reaction concentrations were as follows: 75 μM *holo*-ACP, 1 mM TCEP-HCl, 15 μM KS-CLF, 1.5 μM ScFabD, 10% (v/v) glycerol, and 2 mM ^12/13^C_3_-malonyl-CoA in a total volume of 300 μL. Once all components were combined, reactions were incubated at 30 °C for 16-18 hours, The reaction was extracted (2 × 1 mL) with 99:1 EtOAc:AcOH. The extracts were combined, dried, resuspended in 300 μL MeOH, and analyzed by High-Res LC-MS (see below).

For large-scale reactions (100 × 1 mL or larger), malonyl-CoA was produced *in situ*, for a more cost-effective alternative to commercially available malonyl-CoA. Reactant stocks were prepared in the reaction buffer as follows: 150 μM *holo*-gloACP^Q31G/T35L^, 50 mM TCEP-HCl, 54 μM KS-CLF, 28 μM ScFabD, 309 μM malonyl-CoA synthetase (MatB, from *Rhizobium trifolii*), 250 mM MgCl_2_, 1 M sodium malonate (dibasic monohydrate from Sigma-Aldrich), 250 mM CoA, and 300 mM ATP. Reactions were set up in 100 × 1-dram clear glass vials in 1 mL volumes.

First, *holo*-ACP (13.3 mL), TCEP-HCl (2 mL) and reaction buffer (34.6 mL) were combined in one conical tube and allowed to incubate for 30 °C for 30 min without shaking prior to aliquoting 500 μL into each of 100 × 1-dram clear glass vials. Next, in a second conical tube KS-CLF (18.5 mL), ScFabD (3.6 mL), MatB (6.5 mL), MgCl_2_ (2.8 mL) were combined prior to adding 314 μL into each reaction vial. This was followed by the addition of sodium malonate (100 μL), CoA (20 μL), and finally ATP (67 μL) into each of 100 reaction vials. Each 1-mL reaction was immediately mixed via pipetting. Final concentrations were as follows: 20 μM *holo*-ACP, 1 mM TCEP-HCl, 10 μM KS-CLF, 1 μM ScFabD, 20 μM MatB, 7 mM MgCl_2_, 100 mM sodium malonate, 5 mM CoA, and 20 mM ATP. All 100 × 1 mL reactions were incubated for 18 hours at 30 °C, 80 rpm. Each 1-mL reaction was then extracted 3 × 5 mL of EtOAc:AcOH (99:1). Organic extracts were combined and dried under vacuum, and residual AcOH was removed using toluene azeotrope. Extracts were analyzed by High-Res LC-MS (see below).

### gloPKS *in vitro* reactions with salicyl priming

*In vitro* reaction protocols were adapted from Cheng *et al*.^21^ and run in 50 mM sodium phosphate buffer, pH 7.6. In brief, *holo*-gloACP^Q31G/T35L^ and TCEP-HCl were incubated for 30 minutes at 30 °C followed by the sequential addition of gloSCL, sodium salicylate, MgCl_2_, and ATP. After an additional 30 min incubation at 30 °C, ScFabD, gloKS-CLF, glycerol, and ^12/13^C_3_-malonyl-CoA were added and all components were incubated for 3 hours at 30 °C. Final reaction concentrations were as follows: 75 μM *holo*-gloACP, 1 mM TCEP-HCl, 1.5 μM gloSCL, 0.5 mM sodium salicylate, 5 mM MgCl_2_, 5 mM ATP, 15 μM gloKS-CLF, 1.5 μM FabD, 10% (v/v) glycerol, and 2 mM ^12/13^C_3_-malonyl-CoA in a total volume of 300 μL. The reaction was extracted (2 × 1 mL) of 99:1 EtOAc:AcOH. The extracts were combined, dried, resuspended in 300 μL MeOH, and analyzed by High-Res LC-MS (see below).

### Characterization of *in vitro* reaction products by High-Resolution LC-MS

Molecules were analyzed on a Waters Acquity I-Class UPLC system coupled to a Synapt G2Si HDMS mass spectrometer in positive ion mode with a heated electrospray ionization (ESI) source in a Z-spray configuration. For small molecules, LC separation was performed on a Waters Acquity UPLC BEH C18 (130 Å, 2.1 mm x 50 mm packed with 1.7 μm C18 particles) using an 0.6 mL/min gradient of 95/5 to 15/85 A/B over the course of four minutes. Eluent A is 0.1% (v/v) formic acid in water and B is 0.1% (v/v) formic acid in acetonitrile. Conditions on the mass spectrometer were as follows: capillary voltage 0.5 kV, sampling cone 40 V, source offset 80 V, source 120 °C, desolvation 250 °C, cone gas 0 L/h, desolvation gas 1000 L/h and nebulizer 6.5 bar. The analyzer was operated in resolution mode and low energy data was collected between 100 and 2000 Da at 0.2 sec scan time. Data was collected using a 20–40 V ramp trap collision energy. Masses were extracted from the TOF MS TICs using a 0.01 or 0.05 Da abs width.

## Supplementary Information

The following Supporting Information is available free of charge at the ACS website.

### Detailed Methods

Table S1. Homology comparison of thermorubin-producing proteins to *Gloeocapsa* homologs. Table S2. Plasmid, primer, and amino acid sequences of all proteins used in this study.

Table S3. List of experimental molecular weights of the ACPs obtained by deconvoluting LC-MS results.

Figure S1. *Gloeocapsa* sp. PCC 7428 type II polyketide synthase biosynthetic gene cluster organization.

Figure S2. SDS-PAGE of heterologously expressed and purified core *Gloeocapsa* PKS proteins.

Figure S3. Tandem proteolysis analysis of gloKS-CLF.

Figure S4. LC-MS spectrum gloACP heterologously expressed in *E. coli* BAP1 cells, resulting in *apo*-gloACP.

Figure S5. LC-MS spectrum of gloACP reacted with Sfp (BAP1 and *in vitro*), resulting in *apo*-gloACP.

Figure S6. LC-MS spectrum of gloACP coexpressed with gloPPT, resulting in *apo*-gloACP.

Figure S7. LC-MS spectrum of gloACP coexpressed with gloPPT with coenzyme A, DTT and MgCl_2_, resulting in *apo*-gloACP.

Figure S8. Western Blotting of His_6_-gloPPT.

Figure S9. LC-MS spectrum of gloACP reacted *in vitro* with gloPPT with coenzyme A, DTT and MgCl_2_, resulting in *apo*-gloACP.

Figure S10. LC-MS spectrum of gloACP coexpressed & copurified with gloPPT, followed by incubation with gloSCL, coenzyme A, DTT and MgCl_2_, resulting in *apo*-gloACP.

Figure S11. Backbone NMR assignment and overlay of the [^1^H, ^15^N]-HSQC NMR spectra of ^13^C, ^15^N-labeled wild-type gloACP and gloACP^Q31G/T37L^.

Figure S12. CS-Rosetta convergence plot for gloACP.

Figure S13. [^1^H, ^15^N]-HSQC and 1D ^1^H NMR spectra of ^13^C, ^15^N-labeled *apo*-gloACP (wild-type and Q31G/T35L) upon titration with unlabeled Sfp.

Figure S14. Multiple sequence alignment of various CPs and non-actinobacterial ACPs with varying Sfp-compatibility.

Figure S15. LC-MS spectrum of purified *apo*-gloACP^Q31G/T35L^.

Figure S16. LC-MS spectrum of gloACP^Q31G/T35L^ reacted with Sfp via heterologous expression in *E. coli* BAP1, resulting in *holo*-gloACP^Q31G/T35L^.

Figure S17. LC-MS spectrum of gloACP^Q31G^ reacted with Sfp (BAP1 and *in vitro*), resulting in *apo*-gloACP^Q31G^.

Figure S18. LC-MS spectrum of gloACP^T35L^ reacted with Sfp (BAP1 and *in vitro*), resulting in minimal *holo*-gloACP^T35L^.

Figure S19. LC-MS spectrum of gloACP^Q31G/T35I^ reacted with Sfp (BAP1 and *in vitro*), resulting in *holo*-gloACP^Q31G/T35I^.

Figure S20. LC-MS spectrum of purified *apo*-dacACP.

Figure S21. LC-MS spectrum of dacACP reacted with Sfp (BAP1 and *in vitro*), resulting in minimal *holo*-dacACP.

Figure S22. LC-MS spectrum of purified *apo*-panACP.

Figure S23. LC-MS spectrum of panACP reacted with Sfp (BAP1 and *in vitro*), resulting in *apo*-panACP.

Figure S24. LC-MS spectrum of purified *apo*-dacACP^T43L^.

Figure S25. LC-MS spectrum of dacACP^T43L^ reacted with Sfp (BAP1 and *in vitro*), resulting in *holo*-dacACP^T43L^.

Figure S26. LC-MS spectrum of purified *apo*-panACP^A30G/T34L/A37V^.

Figure S27. LC-MS spectrum of panACP^A30G/T34L/A37V^ reacted with Sfp (BAP1 and *in vitro*), resulting in *holo*-panACP^A30G/T34L/A37V^.

Figure S28. LC-MS spectrum of malonyl-gloACP^Q31G/T35L^ from self-malonylation of *holo*-gloACP^Q31G/T35L^ upon incubation with malonyl CoA.

Figure S29. LC-MS spectrum of malonyl-gloACP^Q31G/T35L^ catalyzed by ScFabD. Figure S30. LC-MS spectrum of malonyl-gloACP^Q31G/T35L^ catalyzed by EcFabD.

Figure S31. Full extracted high resolution ion chromatograms of product peaks of *m/z* = 451, 469, 475, 493, and 511 produced by the reconstituted core gloPKS in the absence of gloSCL/ salicylate.

Figure S32. LC-MS spectrum of salicyl-gloACP^Q31G/T35L^ primed by gloSCL-facilitated loading of salicylic acid.

Figure S33. Full extracted high resolution ion chromatograms of product peaks of *m/z* = 529 and 547 produced by the reconstituted core gloPKS in the presence of gloSCL/ salicylate.

## Acknowledgements/Funding

We thank the Wistar Institute for conducting tandem proteolysis experiments and Professor Dominic Campopiano (University of Edinburgh) for providing the FabD expression plasmid. We are thankful to the Research Corporation for Scientific Advancement (Cottrell Scholars Collaborative Catalyzing Joint Research grant) for funding and supporting the collaboration between LKC and UAH. Generous support from the National Science Foundation (CAREER CHE-1652424 and CHE-2201984) and the Henry Dreyfus Teacher Scholar Award (TH-10-2020) to LKC is acknowledged. UAH acknowledges support by the Center of Biomolecular Magnetic Resonance (BMRZ) funded by the state of Hesse and the Fulbright-Cottrell Award (funded by the German-American Fulbright commission, the Research Corporation for Science

Advancement (RCSA) and the Bundesministerium für Bildung und Forschung (BMBF)). This work was supported by the Deutsche Forschungsgemeinschaft (DFG, German Research Foundation) through the collaborative research center SFB1127/3 ChemBioSys, project ID 3974852 (to PS and UAH) and the Cluster of Excellence “Balance of the Microverse” EXC 2051 – Project-ID 390713860 (to UAH). UAH acknowledges an instrumentation grant for a high-field NMR spectrometer by the REACT-EU EFRE Thuringia (Recovery assistance for cohesion and the territories of Europe, European Fonds for Regional Development, Thuringia) initiative of the European Union. LKC and UAH would like to acknowledge S. Ronco, the entire Cottrell community and A. Cabana for constant encouragement.

## TOC Graphic

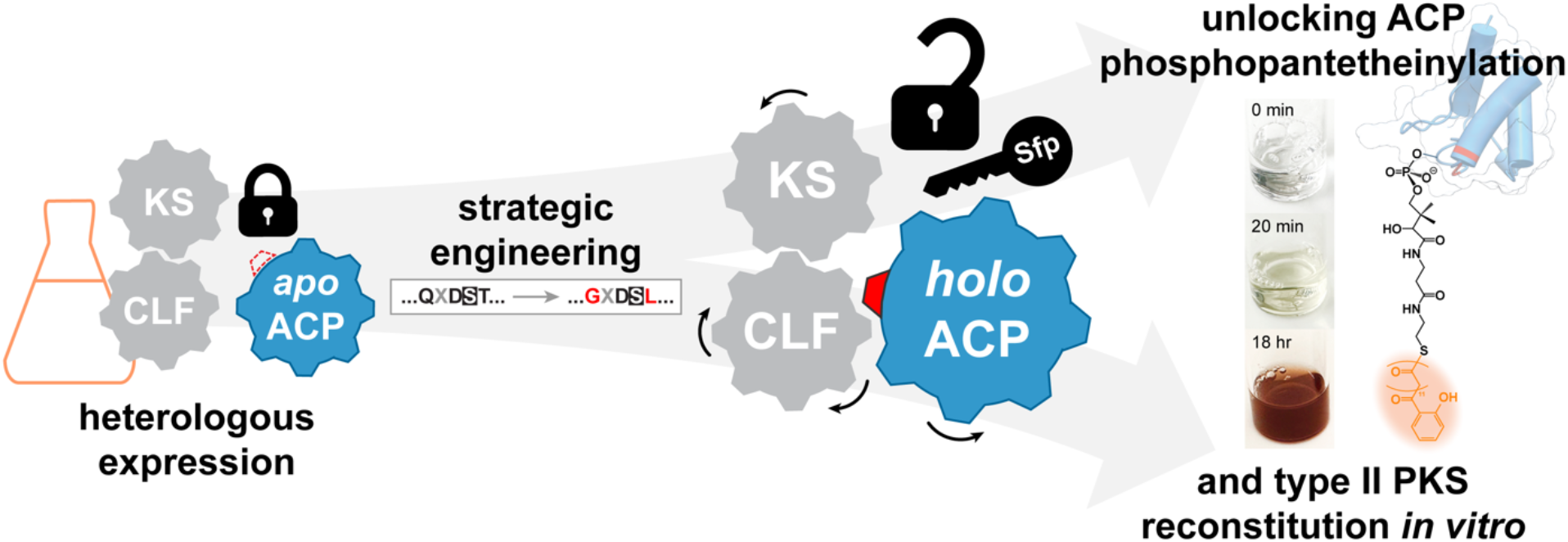

