## Supporting Information for "Strategic acyl carrier protein engineering enables functional type II polyketide synthase reconstitution *in vitro*"

| **Table of Contents** | **Page** |
|---|---|
| **Detailed Methods** | 3 |
| **Table S1.** Homology comparison of thermorubin-producing proteins to *Gloeocapsa* homologs. | 4 |
| **Table S2.** Plasmid, primer, and amino acid sequences of all proteins used in this study. | 5–8 |
| **Table S3.** List of experimental molecular weights of the ACPs obtained by deconvoluting LC-MS results. | 9 |
| **Figure S1.** *Gloeocapsa* sp. PCC 7428 type II polyketide synthase biosynthetic gene cluster organization. | 10 |
| **Figure S2.** SDS-PAGE of heterologously expressed and purified core *Gloeocapsa* PKS proteins. | 11 |
| **Figure S3.** Tandem proteolysis analysis of gloKS-CLF. | 12 |
| **Figure S4.** LC-MS spectrum gloACP heterologously expressed in *E. coli* BAP1 cells, resulting in *apo*-gloACP. | 13 |
| **Figure S5.** LC-MS spectrum of gloACP reacted with Sfp (BAP1 and *in vitro*), resulting in *apo*-gloACP. | 14 |
| **Figure S6.** LC-MS spectrum of gloACP coexpressed with gloPPT, resulting in *apo*-gloACP. | 15 |
| **Figure S7.** LC-MS spectrum of gloACP coexpressed with gloPPT with coenzyme A, DTT and $MgCl_2$, resulting in *apo*-gloACP. | 16 |
| **Figure S8.** Western Blotting of $His_6$-gloPPT. | 17 |
| **Figure S9.** LC-MS spectrum of gloACP reacted *in vitro* with gloPPT with coenzyme A, DTT and $MgCl_2$, resulting in *apo*-gloACP. | 18 |

**Figure S10.** LC-MS spectrum of gloACP coexpressed & copurified with gloPPT, followed by incubation with gloSCL, coenzyme A, DTT and $MgCl_2$, resulting in *apo*-gloACP. 19
**Figure S11.** Backbone NMR assignment and overlay of the [$^1H$, $^{15}N$]-HSQC NMR spectra of $^{13}C$, $^{15}N$-labeled wild-type gloACP and gloACP$^{Q31G/T37L}$. 20
**Figure S12.** CS-Rosetta convergence plot for gloACP. 21
**Figure S13.** [$^1H$, $^{15}N$]-HSQC and 1D $^1H$ NMR spectra of $^{13}C$, $^{15}N$-labeled *apo*-gloACP (wild-type and Q31G/T35L) upon titration with unlabeled Sfp. 22
**Figure S14.** Multiple sequence alignment of various CPs and non-actinobacterial ACPs with varying Sfp-compatibility. 23
**Figure S15.** LC-MS spectrum of purified *apo*-gloACP$^{Q31G/T35L}$. 24
**Figure S16.** LC-MS spectrum of gloACP$^{Q31G/T35L}$ reacted with Sfp via heterologous expression in *E. coli* BAP1, resulting in *holo*-gloACP$^{Q31G/T35L}$. 25
**Figure S17.** LC-MS spectrum of gloACP$^{Q31G}$ reacted with Sfp (BAP1 and *in vitro*), resulting in *apo*-gloACP$^{Q31G}$. 26
**Figure S18.** LC-MS spectrum of gloACP$^{T35L}$ reacted with Sfp (BAP1 and *in vitro*), resulting in minimal *holo*-gloACP$^{T35L}$. 27
**Figure S19.** LC-MS spectrum of gloACP$^{Q31G/T35I}$ reacted with Sfp (BAP1 and *in vitro*), resulting in *holo*-gloACP$^{Q31G/T35I}$. 28
**Figure S20.** LC-MS spectrum of purified *apo*-dacACP. 29
**Figure S21.** LC-MS spectrum of dacACP reacted with Sfp (BAP1 and *in vitro*), resulting in minimal *holo*-dacACP. 30
**Figure S22.** LC-MS spectrum of purified *apo*-panACP. 31
**Figure S23.** LC-MS spectrum of panACP reacted with Sfp (BAP1 and *in vitro*), resulting in *apo*-panACP. 32
**Figure S24.** LC-MS spectrum of purified *apo*-dacACP$^{T43L}$. 33
**Figure S25.** LC-MS spectrum of dacACP$^{T43L}$ reacted with Sfp (BAP1 and *in vitro*), resulting in *holo*-dacACP$^{T43L}$. 34
**Figure S26.** LC-MS spectrum of purified *apo*-panACP$^{A30G/T34L/A37V}$. 35
**Figure S27.** LC-MS spectrum of panACP$^{A30G/T34L/A37V}$ reacted with Sfp (BAP1 and *in vitro*), resulting in *holo*-panACP$^{A30G/T34L/A37V}$. 36
**Figure S28.** LC-MS spectrum of malonyl-gloACP$^{Q31G/T35L}$ from self-malonylation of *holo*-gloACP$^{Q31G/T35L}$ upon incubation with malonyl CoA. 37
**Figure S29.** LC-MS spectrum of malonyl-gloACP$^{Q31G/T35L}$ catalyzed by ScFabD. 38
**Figure S30.** LC-MS spectrum of malonyl-gloACP$^{Q31G/T35L}$ catalyzed by EcFabD. 39
**Figure S31.** Full extracted high resolution ion chromatograms of product peaks of *m/z* = 451, 469, 475, 493, and 511 produced by the reconstituted core gloPKS in the absence of gloSCL/ salicylate. 40
**Figure S32.** LC-MS spectrum of salicyl-gloACP$^{Q31G/T35L}$ primed by gloSCL-facilitated loading of salicylic acid. 41
**Figure S33.** Full extracted high resolution ion chromatograms of product peaks of *m/z* = 529 and 547 produced by the reconstituted core gloPKS in the presence of gloSCL/ salicylate. 42

### Detailed Methods

**Coexpression of gloACP and gloPPT.**
pGloPPTase (sequence can be found in Table S2) was amplified with primers 5'-CTGCAGGCGCGCCGAGCTCGAATTCTCATACGTGTTCGCCC-3' and 5'-ACTTTAATAAGGAGATATACCATGGGCAGCAGCCATCATC-3' and inserted between NcoI and EcoRI restriction sites of digested CDF_Duet1 vector to yield pGloPPT_CDF. *E. coli* BL21(DE3) cells were co-transformed with pGloACP and pGloPPT_CDF. Expression and purification were carried out as outlined in the Methods section with the following exceptions: media was supplemented with 15 µg/mL Kan and 25 µg/mL Strep, and cultures were induced with 500 µM IPTG.

**Western blot.**
Western blotting was used to verify successful *in vitro* isolation of $His_6$-gloPPT. Loading dye was prepared using 2X Laemmli sample buffer with a final concentration of 5% β-mercaptoethanol (βME). Samples were run using a 4–20% Mini-PROTEAN TGX precast protein gel, which was subsequently transferred overnight to a 0.45 mm nitrocellulose membrane using a Mini Trans-Blot Electrophoretic Transfer Cell from BioRad following manufacture protocols. After the overnight transfer, the nitrocellulose membrane was stained with Ponceau S Stain (Sigma Aldrich, Cat. # 6226-79-5) to assess protein transfer success. Afterwards, the membrane was washed multiple times with transfer buffer (25 mM Tris-base, 192 mM glycine, pH 8.3, 20% (v/v) methanol) and TBST buffer (50 mM Tris-HCl, 150 mM NaCl, pH 7.4, 0.1% (v/v) Tween-20), before blocking with TBST buffer + 5% (w/v) Blotto non-fat milk powder. After 1 hour of blocking, the membrane was further rinsed with TBST buffer and stained with 6X-His Tag monoclonal antibody Alexa Fluor™ 488 (Thermo Fisher Scientific) overnight following manufacture protocols and visualized using a FluoroChem M imager (Bio-Techne) (Supplementary Fig. S8).

### Tables and Figures

**Table S1.** Homology comparison between thermorubin-producing proteins and *Gloeocapsa* sp. PCC 7428 PKS homologs. Deduced protein functions of genes in the gloPKS BGC include those essential for *in vitro* biosynthetic functionality (asterisk). N/A indicates no type II polyketide BGC protein in the thermorubin-producing PKS is homologous to the corresponding *Gloeocapsa* protein.

| Protein | Deduced function | Coverage/Percent identity of genes to *L. sacchari* type II PKS homologs |
|---|---|---|
| gloA* = gloKS* | Ketosynthase | 98/66.27 (TheE) |
| gloB* = gloCLF* | Chain length factor | 98/60.86 (TheF) |
| gloC* = gloACP* | Acyl carrier protein | 91/53.42 (family protein) |
| gloD* = gloPPT* | Phosophopantetheinyl transferase | 88/44.26 (superfamily protein) |
| gloE* = gloSCL* | Salicylate CoA ligase | 97/53.77 (TheJ) |
| gloF | Cyclase | 84/54.05 (TheK) |
| gloG | Carboxymuconolactone decarboxylase | 86/64.89 (family protein) |
| gloH | Cyclase | 99/67.27 (TheM) |
| gloI | Cyclase | 90/65.52 (TheN) |
| gloJ | Salicylate synthase | 97/55.24 (TheO) |
| gloK | Proofreading thioesterase | 94/44.09 |
| gloL | Putative 2-dehydropantoate 2-reductase | N/A |
| gloM | Glycosyltransferase | N/A |
| gloN | Hypothetical protein | N/A |
| gloO | O-antigen family ligase protein | N/A |
| gloP | Glycosyltransferase | N/A |
| gloQ | Glycosyltransferase | N/A |
| gloR | Glycosyltransferase | N/A |
| gloS | Glycosyltransferase | N/A |
| gloT | Lipopolysaccharide biosynthesis protein | N/A |

**Table S2.** Primers used for cloning, amino acid sequences, and theoretical MW of proteins used in this study (note: pET28a leader sequence underlined).

| Protein | Plasmid name, Primers (5'→3') &<br>Amino Acid Sequence (confirmed by sequencing) | Theoretical MW (Da) |
|---|---|---|
| *apo*-gloACP<br>(WT) | Plasmid: pGloACP<br><br>Forward primer:<br>CCTGGTGCCGCGCGGCAGCCATATGGTCATGGATGCGCTAAAAG<br>Reverse primer:<br>AAGCTTGTCGACGGAGCTCGAATTCTCATGTCGCCGTAGCCGTA<br><br>AA sequence of expressed protein:<br><u>MGSSHHHHHHSSGLVPRGSH</u>MVMDALKDILVDLGIPEQEITETALLRK<br>DLQLDSTETVDISLGLKRRFGVNVKLESRKDMTLKDVCEMVNSAIAATA<br>TAT | 10957.55 |
| *apo*-gloACP<br>(Q31G/T35L) | Plasmid: pSDM3 (template plasmid = pGloACP)<br><br>Forward primer:<br>CTCACTGGAAACCGTCGATATTTCCC<br>Reverse primer:<br>TCGAGACCCAAGTCTTTGCGCAGCAG<br><br>AA sequence of expressed protein:<br><u>MGSSHHHHHHSSGLVPRGSH</u>MVMDALKDILVDLGIPEQEITETALLRK<br>DLGLDSLETVDISLGLKRRFGVNVKLESRKDMTLKDVCEMVNSAIAATA<br>TAT | 10898.52 |
| *apo*-gloACP<br>(Q31G/T35I) | Plasmid: pSDM4 (template plasmid = pGloACP)<br><br>Forward primer: 5' CTCAATCGAAACCGTCGATATTTCCC 3'<br>Reverse primer: 5' TCGAGACCCAAGTCTTTGCGCAGCAG 3'<br><br>AA sequence of expressed protein:<br><u>MGSSHHHHHHSSGLVPRGSH</u>MVMDALKDILVDLGIPEQEITETALLRK<br>DLGLDSIETVDISLGLKRRFGVNVKLESRKDMTLKDVCEMVNSAIAATA<br>TAT | 10898.52 |
| *apo*-gloACP<br>(Q31G) | Plasmid: pSDM5 (template plasmid = pGloACP)<br><br>Forward primer: 5' CAAAGACTTGGGTCTCGACTCAACAGAAACCG 3'<br>Reverse primer: 5' CGCAGCAGTGCTGTTTCT 3'<br><br>AA sequence of expressed protein:<br><u>MGSSHHHHHHSSGLVPRGSH</u>MVMDALKDILVDLGIPEQEITETALLRK<br>DLGLDSTETVDISLGLKRRFGVNVKLESRKDMTLKDVCEMVNSAIAATA<br>TAT | 10886.47 |
| *apo*-gloACP<br>(T35L) | Plasmid: pSDM6 (template plasmid = pGloACP)<br><br>Forward primer: 5' ACTCGACTCATTAGAAACCGTCGATATTTCC 3'<br>Reverse primer: 5' TGCAAGTCTTTGCGCAGC 3'<br><br>AA sequence of expressed protein:<br><u>MGSSHHHHHHSSGLVPRGSH</u>MVMDALKDILVDLGIPEQEITETALLRK<br>DLQLDSLETVDISLGLKRRFGVNVKLESRKDMTLKDVCEMVNSAIAATA<br>TAT | 10969.60 |
| gloKS-CLF | plasmid: pGloKSCLF<br><br>Forward primer:<br>CCTGGTGCCGCGCGGCAGCCATATGAAACGAGTAGTCATTACAG | $His_6$ KS:<br>47432.43 |

| | | |
|---|---|---|
| | Reverse primer:<br>AAGCTTGTCGACGGAGCTCGAATTCTCACTTAGCAATCACAAAC<br><br>AA sequence of expressed protein<br>His6 KS:<br>MGSSHHHHHHSSGLVPRGSHMKRVVITGIGVVAPLGIGKEQFWKNAIR<br>GQSYLQADPEMEAMGIKSKVVCRVADFDLSDYCSGAEFDHLVEQDRV<br>VQFGVVSGTAAIADSGLDLSQEDPESLGIIFSSAIGGTPTIQKIFERCSEK<br>GTQPLKHVATGENFYNAGMFNYPAALLARKHGFQGPCTSLSTGCTAG<br>LDALGLSFELIRSGECKVMLAGASEAPLTSLTYATLDVIGSLSVADCEPE<br>KASRPFDAKRGGFVISEAGAVLVLEELEHALNRNAHIYAEVVSYYSVSN<br>AFHMTDLPDHGVPMAAVMERTLHLGNVEPEELDYINAHGSSTPQNDL<br>FETNAYKQVLGDKAYRLPISSTKSMIGHSLSSASLVGVVATLGAIELSVI<br>HPTANYEFPDPNCDLDYVPNEARSTEVNTALLTASGFGGIHSAAIFKKY<br>QESLGE<br><br>CLF:<br>MSKHDVVITGIGIINPAGIGKDEFWHNISTGKSAIREISRFDSTDFPTKVA<br>GEIAEFEPADYIPRRFIVKTDRFTHYALAATELALQDATLDLTQEDSYRV<br>GVWFGNNAGGWDICERGFYELYNDGATMVNPWQATAWFPTAAQGY<br>VTIRYGIRGYSKSFVCDRASGASGLYFGIKSIQEGFNDVVIAGGSEAPIT<br>RFGMTCYYETGEVSAATDPEKAYLPFDRNRTGLVLGEGSTVLVLESEE<br>HARNRGAKIYGKVVSGCMTTDTDPTSGIHFERCMTRAIQSAQIQPTDID<br>VVLAEGCGTQQSDRIEGEAISTVFAQAPKVAVSVPKALYGHLYGASCV<br>TEVACSLLAMETEQLPTMSQTEPDADCRLNFVTQPQNHPVRHALVNS<br>RAREGVNASFVIAK | CLF:<br>43813.18 |
| gloPPT | Plasmid: pGloPPTase<br><br>Forward primer: GTGCCGCGCGGCAGCCATATG<br>Reverse primer: AAGCTTGTCGACGGAGCTCGAATTC<br><br>AA sequence of expressed protein:<br>MGSSHHHHHHSSGLVPRGSHMMPTGKGIQGFMYLQIAQPRQLAEIKG<br>IGIDIAPVSKIASLVSRYNSETLTLLFTPREIEQCQSAPYPNRYYAVCFAA<br>KEAVGKALGTGLVDINWNEIESIISQSELTIKLRGAAKQKALQCGVKAW<br>LATWCYWDDYVMVHVLSKGEHV | 18717.59 |
| gloSCL | Plasmid: pGloSCL<br><br>Forward primer:<br>AGCGGCCTGGTGCCGCGCGGCAGCCATATGAATCAGTACGAACAA<br>TTAC<br>Reverse primer:<br>GCCGCAAGCTTGTCGACGGAGCTCGAATTCTTATTGAACCAGTTGA<br>CTG<br><br>AA sequence of expressed protein:<br>MGSSHHHHHHSSGLVPRGSHMNQYEQLPDIFNVAAYFIKGNLHKGYG<br>ERIALYHQDDTYTYRKVSNEICAAAGLLAELGLERENRFAILLPDSPDFV<br>FAFWGAIWLGAVPVPINTACNLDDIEYILQDCRAKILLTTQEWQDKLSPI<br>QSPFLRNILLTDGENSFRTLASSFSQELPPAQTSPDEPAFWLYTSGST<br>GRPKGAIHLHRSMVFCAEQYGKATLGLHQDDITYSIAKMPFAYGLGNT<br>LYMPMAVGAASILSDANNAFDIIADIHRHRPTILFAIPATYASILAVQDIAP<br>LDASTLRLCVSAAEQLPKSIWQRWRSTYGIEICEGIGTTEFLHIFLSNRL<br>GECRPGSSGKPVVGYDVRIIDENGVSMPTGEIGNLQVGGDSLMLRYW<br>NRHQETRQVIHGNTMRTGDKYLCDADGYFWFMGRKDDLFKVNGQW<br>VSPFEIEDVLLQHESVLDVAVVPESESGENLTQVVAYISLKAGFSESVE<br>LEESIRRFAKMQLPRFKAPKKIQFLERLPRTSTGKIHRKALLKASQLVQ | 59879.20 |

| | | |
|---|---|---|
| *apo*-panACP<br>(WT) | Plasmid: pPanACP<br><br>Twist Bioscience codon optimized DNA purchased:<br>GTGCCGCGCGGCAGCCATATGATGGTATTCGAGAAGGTCAAAGCG<br>ATTATTGAGGATATTGGGATTGAAGACGAGATCAT<br>TGAGTCGAGCCGCTTGTACGATGACTTAGCGCTTGATTCTACTGAG<br>TTAGCATTAGTATCCACGGCTCTGGCAAAGGCAT<br>TTGGAATCTTTATTGAAAGTCGTGTACTGAAGACCTATTCCGTTGCA<br>CAAGTGATTGAGGCTGTCGCCTTGAAGGCGTGA<br>GAATTCGAGCTCCGTCGACAAGCTT<br><br>Forward primer: GTGCCGCGCGGCAGCCATATG<br>Reverse primer: AAGCTTGTCGACGGAGCTCGAATTC<br><br>AA sequence:<br>MGSSHHHHHHSSGLVPRGSHMMVFEKVKAIIEDIGIEDEIIESSRLYDDL<br>ALDSTELALVSTALAKAFGIFIESRVLKTYSVAQVIEAVALKA | 10169.66 |
| *apo*-panACP<br>(A30G/T34L/A37V) | Plasmid: pPaACP_SDM2 (template plasmid = pPanACP)<br><br>Forward primer: CTGGAGTTAGTATTAGTATCCACGGCTCTG<br>Reverse primer: AGAATCAAGCCCTAAGTCATCGTACAAGCG<br><br>AA sequence:<br>MGSSHHHHHHSSGLVPRGSHMMVFEKVKAIIEDIGIEDEIIESSRLYDDL<br>GLDSLELVLVSTALAKAFGIFIESRVLKTYSVAQVIEAVALKA | 10195.74 |
| *apo*-dacACP<br>(WT) | Plasmid: pDacACP<br><br>Twist Bioscience codon optimized DNA purchased:<br>GTGCCGCGCGGCAGCCATATGATGAAAGATCATGTTTCGATTGAAG<br>CTATTGTTATTAACGCCTTAAAGGCATTGGTCGAGTCTGAGGGGTTA<br>AAAGTTGACATCACTCGCGCTTCTATTATGGCGGACGACTTAGGTG<br>TTGACTCAACTGAACTTGTGTTTATTCTGCTTGAGATTGAGAACCAA<br>ACAGCCCAAGCTCTGAAAGACATCGACTATGGACGTATCAGCACTG<br>TTGGTGATTTAATCGATGCTGCCCAAGAAGCGGCAGGTGTATAGGA<br>ATTCGAGCTCCGTCGACAAGCTT<br><br>Forward primer: GTGCCGCGCGGCAGCCATATG<br>Reverse primer: AAGCTTGTCGACGGAGCTCGAATTC<br><br>AA sequence of expressed protein:<br>MGSSHHHHHHSSGLVPRGSHMMKDHVSIEAIVINALKALVESEGLKVDI<br>TRASIMADDLGVDSTELVFILLEIENQTAQALKDIDYGRISTVGDLIDAAQ<br>EAAGV | 11249.71 |
| *apo*-dacACP<br>(T43L) | Plasmid: pDaACP_SDM1 (template plasmid = pDacACP)<br><br>Forward primer:<br>TGTTGACTCACTGGAACTTGTGTTTATTCTGCTTGAG<br>Reverse primer: CCTAAGTCGTCCGCCATA<br><br>AA sequence of expressed protein:<br>MGSSHHHHHHSSGLVPRGSHMMKDHVSIEAIVINALKALVESEGLKVDI<br>TRASIMADDLGVDSLELVFILLEIENQTAQALKDIDYGRISTVGDLIDAAQ<br>EAAGV | 11261.77 |

| | | |
|---|---|---|
| MatB | Plasmid: pMatB<br><br>Twist Bioscience codon optimized DNA purchased:<br>ATGTCCAATCACCTGTTTGATGCGATGCGAGCTGCAGCACCGGGAA<br>ATGCCCCGTTTATTCGAATTGACAATACACGTACGTGGACGTACGA<br>TGATGCGTTTGCCTTAAGTGGGCGAATCGCGTCTGCAATGGATGCA<br>CTGGGGATCCGTCCAGGAGATCGTGTGGCTGTTCAAGTTGAAAAGT<br>CAGCTGAAGCTCTGATTTTATACTTAGCATGCCTGCGGTCGGGTGC<br>GGTATATTTACCACTTAATACAGCGTACACCTTAGCCGAATTGGACT<br>ACTTCATTGGTGACGCAGAACCCCGGCTGGTTGTGGTGGCTAGCT<br>CAGCAAGAGCCGGTGTCGAAACGATTGCGAAACCACGTGGCGCCA<br>TAGTAGAGACACTGGATGCCGCCGGAAGTGGTAGCCTGCTCGACC<br>TTGCACGTGATGAACCAGCGGATTTCGTTGACGCGTCACGATCAGC<br>TGACGACTTAGCAGCAATTCTGTATACAAGCGGTACAACCGGGAGA<br>AGCAAAGGTGCCATGTTAACACACGGCAATTTGCTGTCAAATGCAC<br>TTACTCTGCGCGACTTCTGGCGTGTAACAGCTGGTGACCGTTTAAT<br>ACACGCTCTGCCTATTTTCCATACCCACGGGCTGTTTGTGGCTACA<br>AATGTGACGCTTCTTGCTGGTGCGAGTATGTTTCTTTTAAGCAAATT<br>TGATCCAGAGGAAATTCTTTCTTTGATGCCTCAAGCCACAATGTTAA<br>TGGGTGTTCCGACATTTTATGTACGATTACTCCAATCGCCCAGACTG<br>GATAAACAGGCCGTAGCTAATATAAGACTGTTTATCAGCGGAAGCG<br>CCCCGCTCCTGGCGGAGACGCACACAGAATTTCAAGCCCGCACAG<br>GCCATGCTATCCTGGAACGTTATGGAATGACCGAGACGAACATGAA<br>TACCTCAAATCCCTACGAAGGTAAGAGAATAGCAGGCACCGTGGGT<br>TTTCCTTTACCCGACGTTACAGTTCGAGTAACGGACCCGGCAACTG<br>GCTTAGCCCTCCCTCCAGAGCAGACAGGGATGATAGAAATTAAAGG<br>ACCCAATGTCTTTAAAGGATACTGGCGTATGCCTGAGAAGACGGCT<br>GCTGAGTTTACGGCAGATGGGTTCTTTATATCTGGAGACTTAGGAA<br>AGATTGATAGAGATGGGTACGTGCATATTGTGGGACGGGGTAAAGA<br>CTTGGTTATATCCGGCGGTTATAATATTTACCCAAAGGAAGTGGAAG<br>GTGAAATTGATCAAATTGAAGGAGTAGTCGAATCAGCGGTCATTGG<br>TGTTCCTCACCCAGACTTTGGCGAGGGAGTCACCGCGGTAGTTGTA<br>AGAAAACCTGGTGCGGCACTGGACGAGAAAGCGATAGTGTCCGCG<br>TTACAAGATCGTCTGGCTCGGTATAAGCAGCCGAAACGAATTATTTT<br>CGCCGAAGATCTCCCCCGTAATACTATGGGCAAAGTGCAAAAGAAT<br>ATTTTGCGCCAACAGTATGCGGACTTGTACACTCGTACTTAG<br><br>AA sequence:<br>MGSSHHHHHHSSGLVPRGSHMMSNHLFDAMRAAAPGNAPFIRIDNTR<br>TWTYDDAFALSGRIASAMDALGIRPGDRVAVQVEKSAEALILYLACLRS<br>GAVYLPLNTAYTLAELDYFIGDAEPRLVVVASSARAGVETIAKPRGAIVE<br>TLDAAGSGSLLDLARDEPADFVDASRSADDLAAILYTSGTTGRSKGAM<br>LTHGNLLSNALTLRDFWRVTAGDRLIHALPIFHTHGLFVATNVTLLAGA<br>SMFLLSKFDPEEILSLMPQATMLMGVPTFYVRLLQSPRLDKQAVANIRL<br>FISGSAPLLAETHTEFQARTGHAILERYGMTETNMNTSNPYEGKRIAGT<br>VGFPLPDVTVRVTDPATGLALPPEQTGMIEIKGPNVFKGYWRMPEKTA<br>AEFTADGFFISGDLGKIDRDGYVHIVGRGKDLVISGGYNIYPKEVEGEID<br>QIEGVVESAVIGVPHPDFGEGVTAVVVRKPGAALDEKAIVSALQDRLAR<br>YKQPKRIIFAEDLPRNTMGKVQKNILRQQYADLYTRT | 56910.01 |

**Table S3.** List of experimental molecular weights of ACPs obtained via deconvoluting LC-MS results using ESIprot.[1] All assignments include the observed the loss of the *N*-terminal methionine.

| Fig. # | | Major set of peaks | | Second set of peaks | | Third set of peaks | |
|---|---|---|---|---|---|---|---|
| | | Deconvoluted MW ± SD | ACP state | Deconvoluted MW ± SD | ACP state | Deconvoluted MW ± SD | ACP state |
| S20 | dacACP | 11117.5 ± 0.3 | *apo* | | | | |
| S21 | dacACP + Sfp | 11117.5 ± 0.7 | *apo* | 11457.8 ± 0.4 | *holo*<br>(but minimal) | | |
| S24 | dacACP$^{T43L}$ | 11128.7 ± 0.6 | *apo* | | | | |
| S25 | dacACP$^{T43L}$ + Sfp | 11469.4 ± 0.7 | *holo* | | | | |
| S22 | panACP | 10037.4 ± 0.4 | *apo* | | | | |
| S23 | panACP + Sfp | 10037.6 ± 0.5 | *apo*<br>(no activation) | 10215.8 ± 0.5 | *apo* + glucon.178<br>(no activation) | | |
| S26 | panACP$^{A30G/T34L/A37V}$ | 10063.0 ± 0.7 | *apo* | 10241.5 ± 0.6 | *apo* + glucon.178 | | |
| S27 | panACP$^{A30G/T34L/A37V}$ + Sfp | 10403.5 ± 0.4 | *holo* | 10581.6 ± 0.5 | *holo* + glucon.178 | | |
| S04 | gloACP | 10825.3 ± 0.3 | *apo* | 11003.1 ± 0.3 | *apo* + glucon.178 | | |
| S05 | gloACP + Sfp | 10825.3 ± 0.3 | *apo*<br>(no activation) | 11003.5 ± 0.6 | *apo* + glucon.178<br>(no activation) | | |
| S09 | gloACP + gloPPT *in vitro* | 10824.8 ± 0.6 | *apo*<br>(no activation) | 11003.1 ± 0.5 | *apo* + glucon.178<br>(no activation) | | |
| S06 | gloACP/gloPPT coexpression | 10824.9 ± 0.5 | *apo*<br>(no activation) | 11003.2 ± 0.4 | *apo* + glucon.178<br>(no activation) | | |
| S07 | gloACP/gloPPT coexpressed<br>+ DTT, CoA, $MgCl_2$ | 10824.7 ± 0.6 | *apo*<br>(no activation) | 11003.0 ± 0.5 | *apo* + glucon.178<br>(no activation) | | |
| S10 | gloACP/gloPPT coexpressed<br>+ *in vitro* gloSCL, DTT, CoA, $MgCl_2$, ATP, salicylic acid | 10824.7 ± 0.6 | *apo*<br>(no activation) | 11003.2 ± 0.4 | *apo* + glucon.178<br>(no activation) | | |
| S15 | gloACP$^{Q31G/T35L}$ | 10766.0 ± 0.6 | *apo* | 10944.6 ± 0.8 | *apo* + glucon.178 | | |
| S16 | gloACP$^{Q31G/T35L}$ + Sfp | 11106.4 ± 0.5 | *holo* | 11144.9 ± 0.8 | *holo* + acetylation | 11283.8 ± 2.4 | *holo* + glucon.178 |
| S19 | gloACP$^{Q31G/T35I}$ + Sfp | 10766.5 ± 0.8 | *apo* | 11106.6 ± 0.8 | *holo* | | |
| S17 | gloACP$^{Q31G}$ + Sfp | 10753.9 ± 0.6 | *apo*<br>(no activation) | 10932.4 ± 0.6 | *apo* + glucon.178<br>(no activation) | | |
| S18 | gloACP$^{T35L}$ + Sfp | 10837.0 ± 0.3 | *apo* | 11015.2 ± 0.4 | *apo* + glucon.178 | 11177.05 ± 0.9 | *holo* |
| S28 | gloACP$^{Q31G/T35L}$ self-malonylation | 11106.1 ± 0.6 | *holo* | 11284.1 ± 0.5 | *holo* + glucon.178 | 11191.6 ± 0.6 | malonyl |
| S29 | gloACP$^{Q31G/T35L}$ + ScFabD malonylation | 11192.6 ± 0.5 | malonyl | 11370.6 ± 0.7 | malonyl + glucon.178 | | |
| S30 | gloACP$^{Q31G/T35L}$ + EcFabD malonylation | 11192.2 ± 0.8 | malonyl | 11369.9 ± 0.7 | malonyl + glucon.178 | | |
| S32 | *holo*-gloACP$^{Q31G/T35L}$<br>+ *in vitro* gloSCL, salicylic acid, ATP, TCEP | 11226.3 ± 0.8 | salicyl | 11403.9 ± 0.4 | salicyl + glucon.178 | | |


(1) Winkler, R. ESIprot: A Universal Tool for Charge State Determination and Molecular Weight Calculation of Proteins from Electrospray Ionization Mass Spectrometry Data. *Rapid Commun. Mass Spectrom.* **2010**, *24* (3), 285–294.

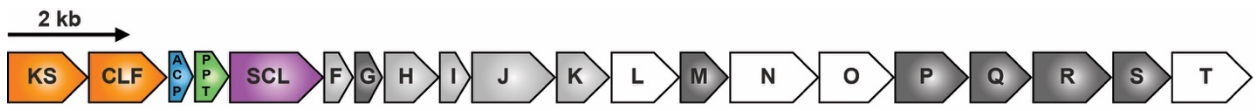


| Protein | Deduced Function | Protein | Deduced Function | Protein | Deduced Function | Protein | Deduced Function |
|---|---|---|---|---|---|---|---|
| gloKS* | Ketosynthase | gloF | Cyclase | gloK | Thioesterase | gloP | Glycosyltransferase |
| gloCLF* | Chain-length factor | gloG | Carboxymuconolactone decarboxylase | gloL | Unknown | gloQ | Glycosyltransferase |
| gloACP* | Acyl carrier protein | gloH | Cyclase | gloM | Glycosyltransferase | gloR | Glycosyltransferase |
| gloPPT* | Phosphopantetheinyl transferase | gloI | Cyclase | gloN | Unknown | gloS | Glycosyltransferase |
| gloSCL* | Salicylate CoA ligase | gloJ | Salicylate synthase | gloO | Unknown | gloT | Unknown |

**Figure S1.** *Gloeocapsa* sp. PCC 7428 type II PKS BGC organization. Deduced gene functions for core PKS enzymes (asterisk), accessory PKS enzymes (light gray), tailoring enzymes (dark gray), and domains of unknown function (white).

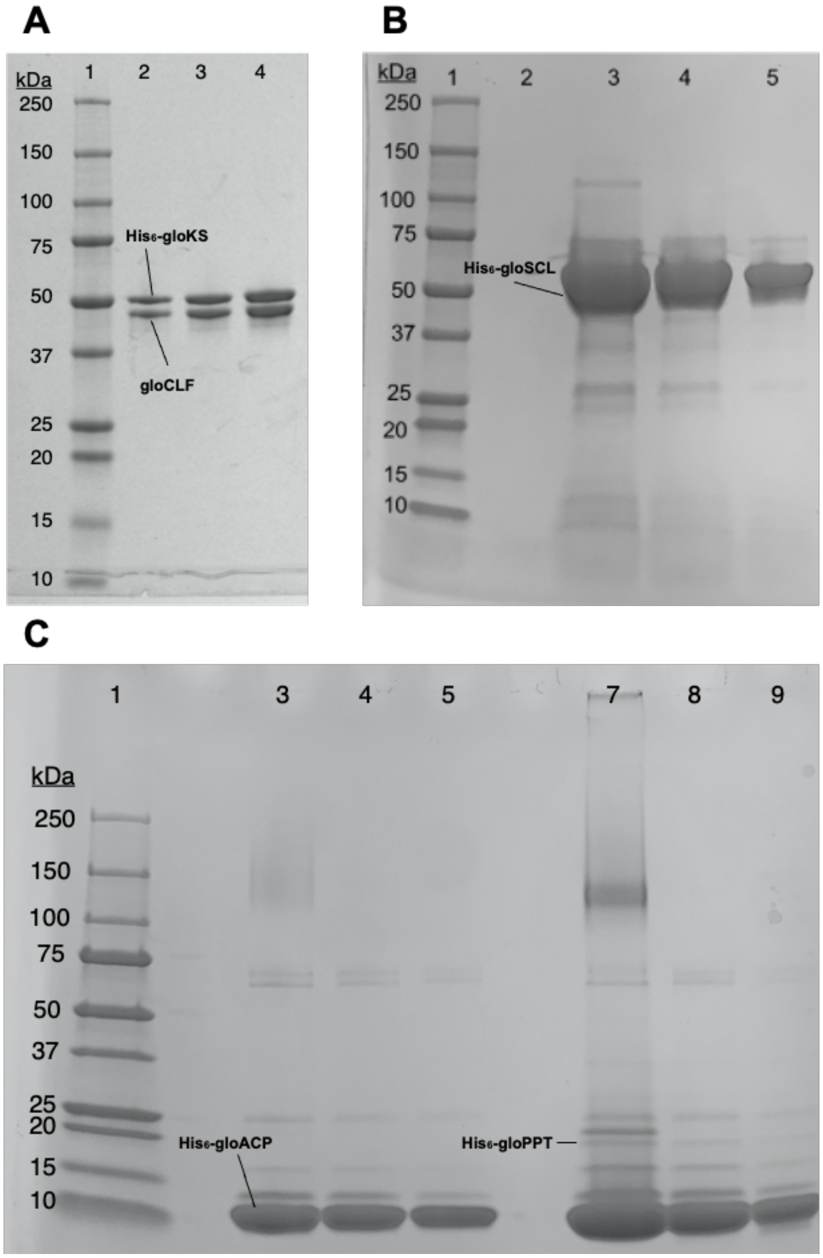


**Figure S2.** SDS-PAGE of heterologously expressed and purified core *Gloeocapsa* sp. PCC 7428 type II PKS (gloPKS) proteins, *i.e.*, gloKS and glCLF, at varying concentrations. Only gloKS carries a $His_6$-tag, but both proteins are expressed under the control of the same promoter and can be purified together. Lane 1 in all gels corresponds to Precision Plus Protein Standards All Blue, BioRad ladder. (A) Ni-NTA purified gloKS-CLF (lanes 2-4). (B) Ni-NTA purified gloSCL (lanes 3-5). (C) Ni-NTA purified $His_6$-gloACP (Lanes 3-5) and co-expressed $His_6$-gloACP and $His_6$-gloPPT (Lanes 7-9).

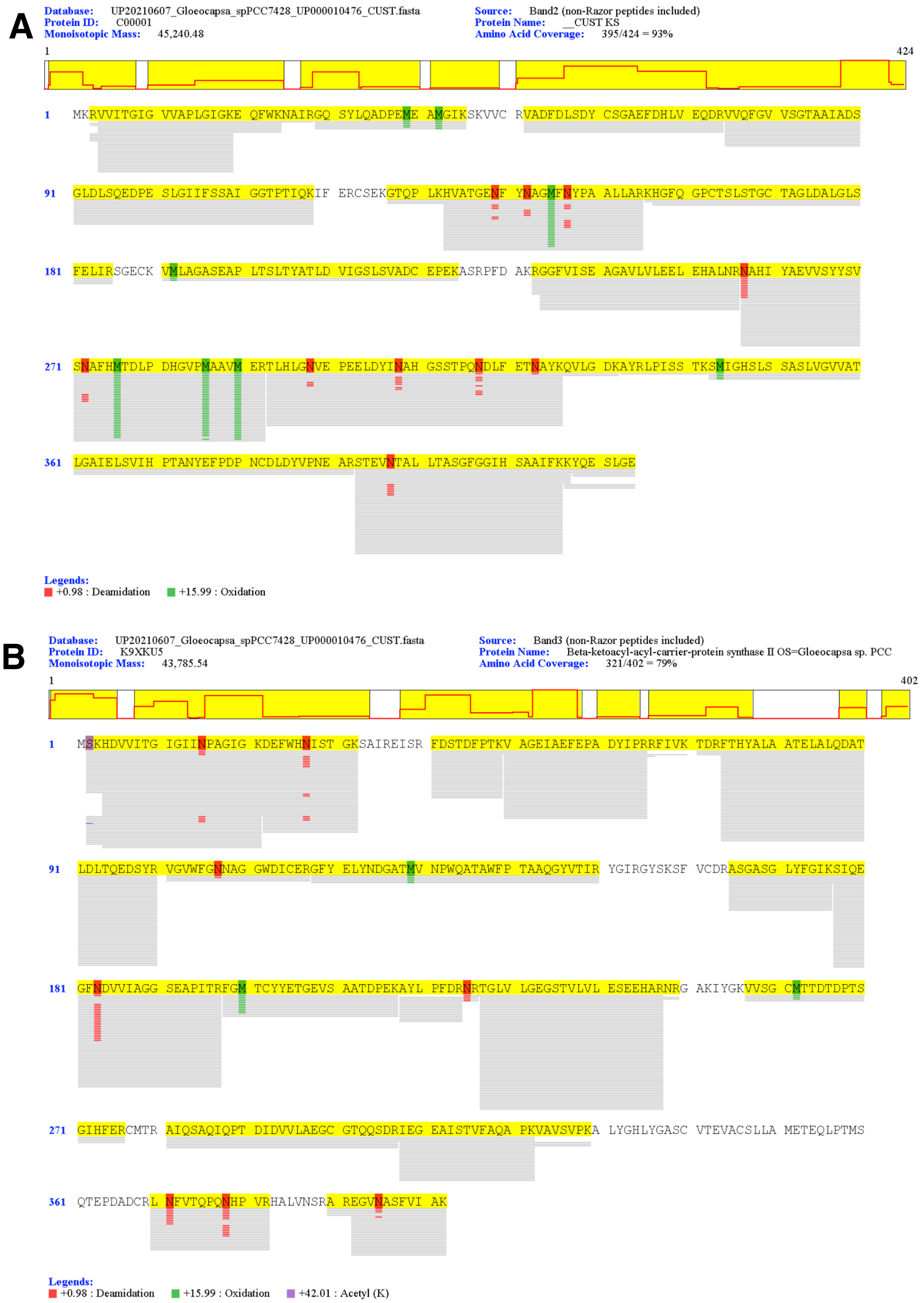

**Figure S3.** Tandem proteolysis analysis of gloKS-CLF. Trypsin digests of ~50 kDa gloKS-CLF band corresponding to $His_6$-gloKS (A) and ~44 kDa gloCLF (B). All expected proteolytic fragments of gloKS could be identified, with grey bar representing frequency of bands observed.

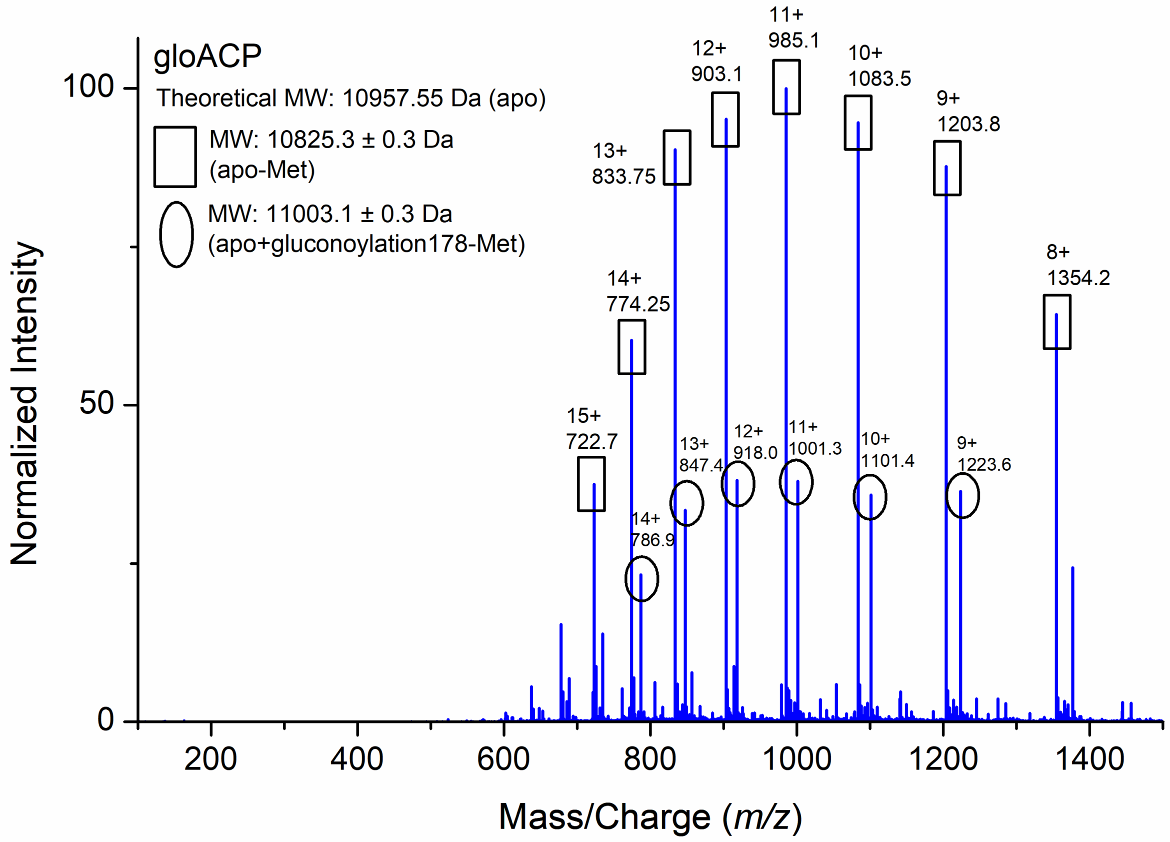


**Figure S4.** LC-MS spectrum of purified gloACP heterologously expressed in *E. coli* BAP1 shows the protein in *apo*-form. No phosphopantetheinylation of ACP was observed.

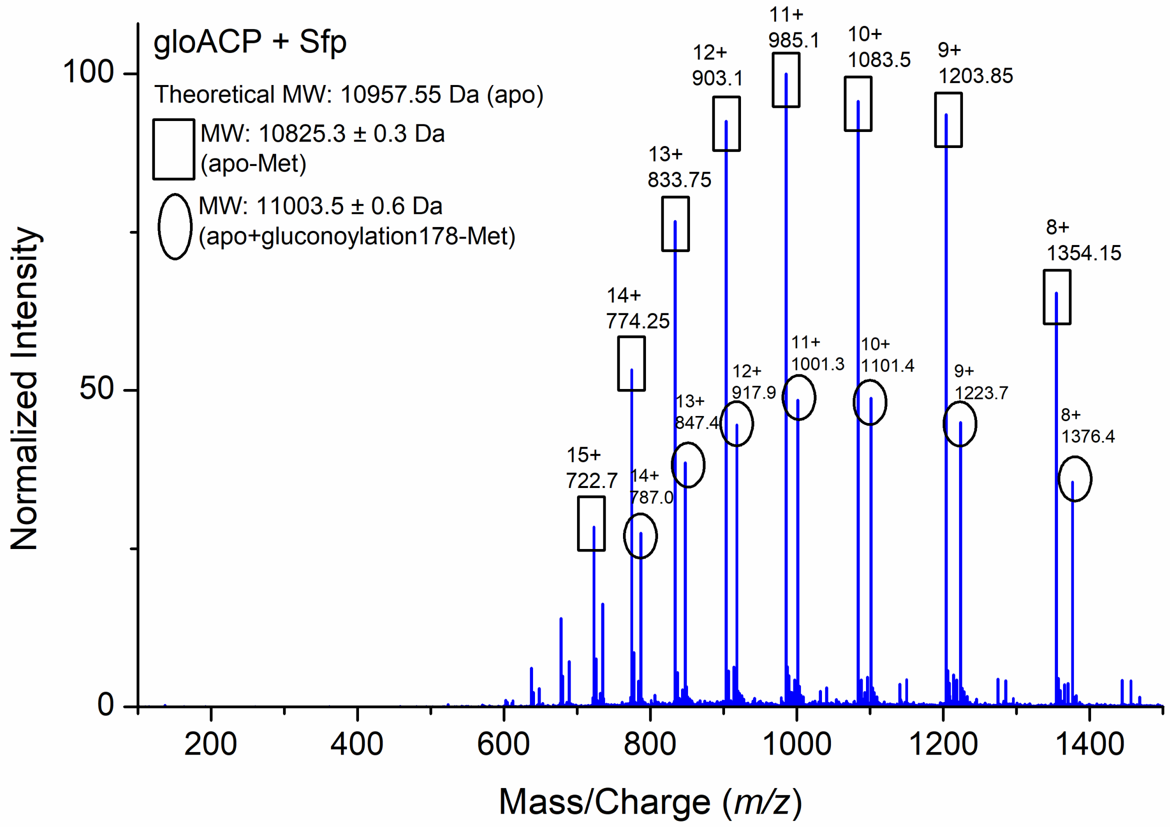


**Figure S5.** LC-MS spectrum of gloACP after incubation with Sfp. GloACP was first expressed and purified from *E. coli* BAP1 and subsequently reacted *in vitro* with Sfp R4-4, DTT, coenzyme A, and $MgCl_2$. No phosphopantetheinylation of ACP was observed.

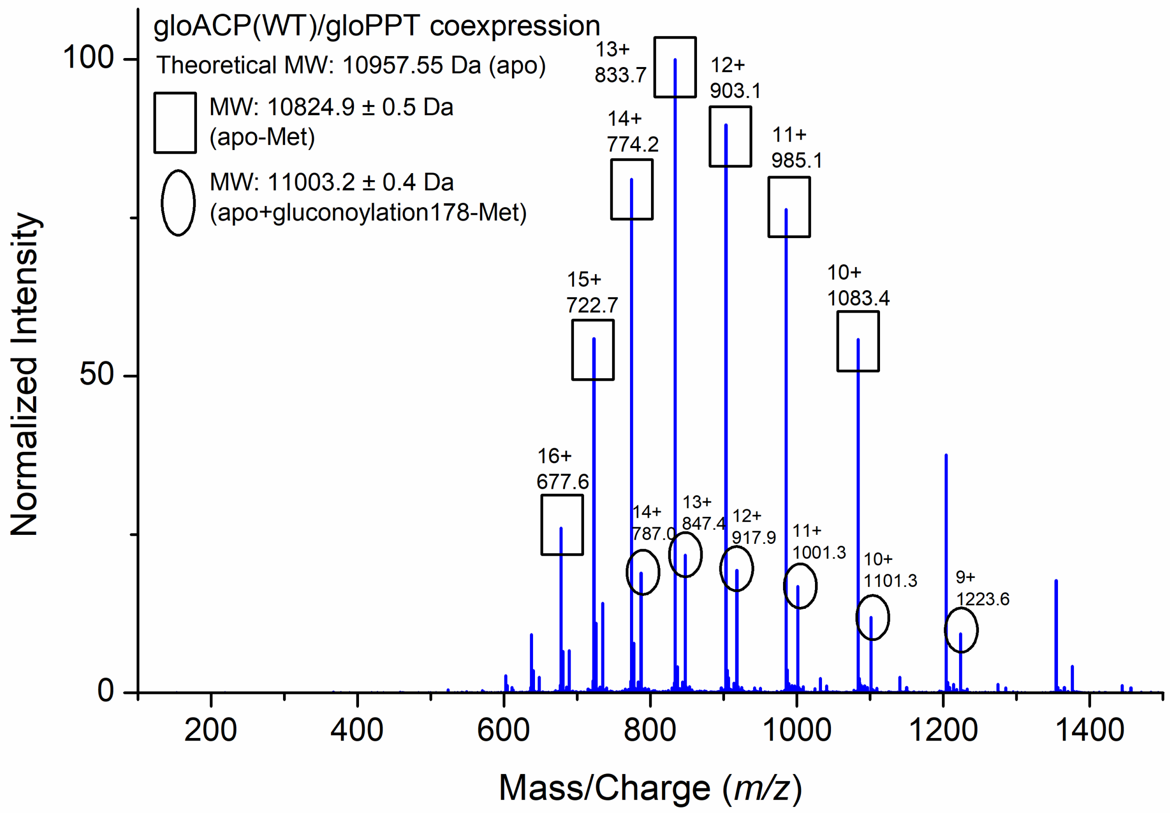


**Figure S6.** LC-MS spectrum of gloACP following heterologous coexpression with gloPPT in *E. coli.* No phosphopantetheinylation of ACP was observed.

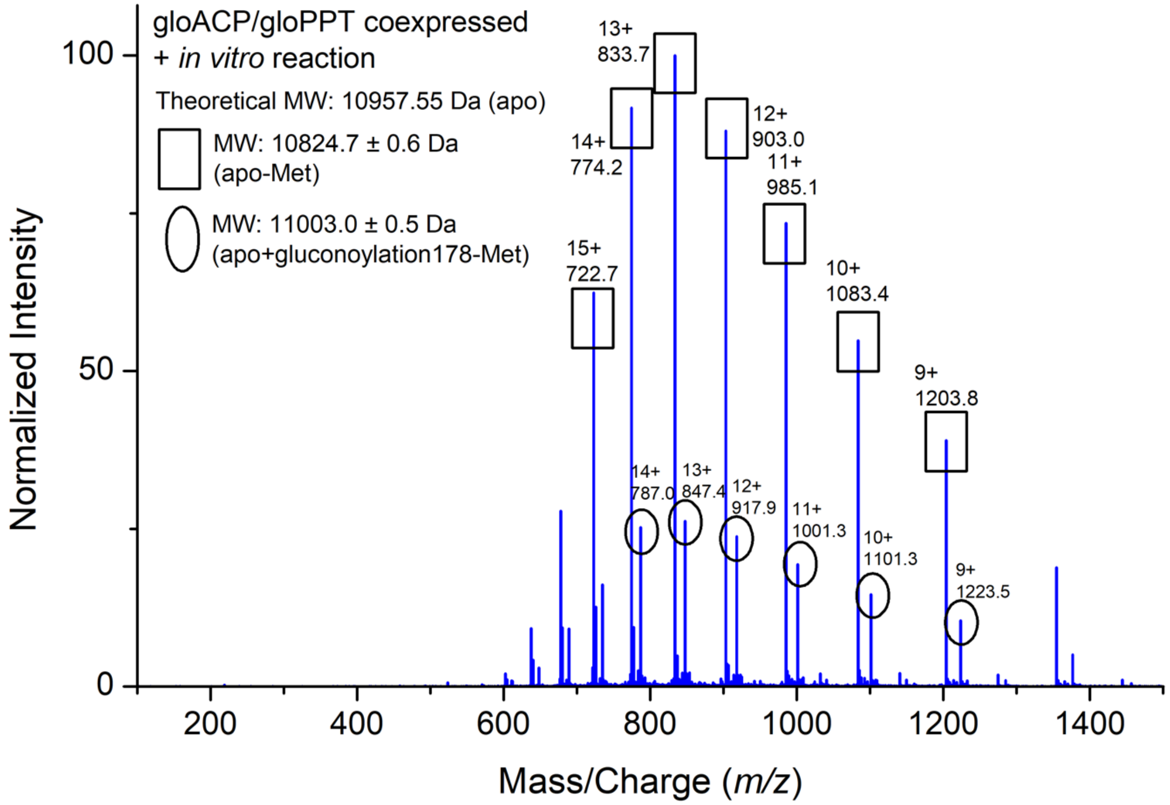


**Figure S7.** LC-MS spectrum of co-expressed/co-purified gloACP and gloPPT Additional reaction components include DTT, coenzyme A, and $MgCl_2$. No phosphopantetheinylation of ACP was observed.

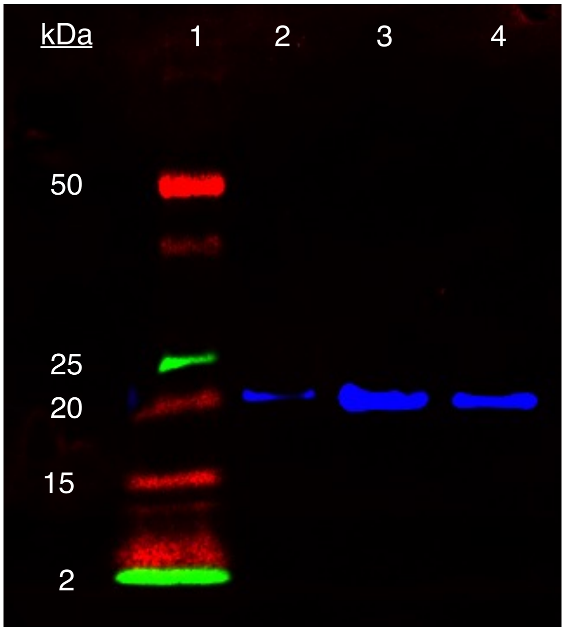


**Figure S8.** Western Blot of His$_6$-gloPPT. Isolation of His$_6$-gloPPT was confirmed by presence of an 18.7 kDa band. His$_6$-tagged gloPPT was labeled using 6X-His Tag monoclonal antibody Alexa Fluor$^{TM}$ 488 overnight and imaged using FluoroChem M imager, showing up as blue bands. Lane 1: ladder (Precision Plus Protein Dual Color Standards, BioRad). Lane 2: post-induction sample of gloPPT. Lanes 3 & 4: post-purification gloPPT.

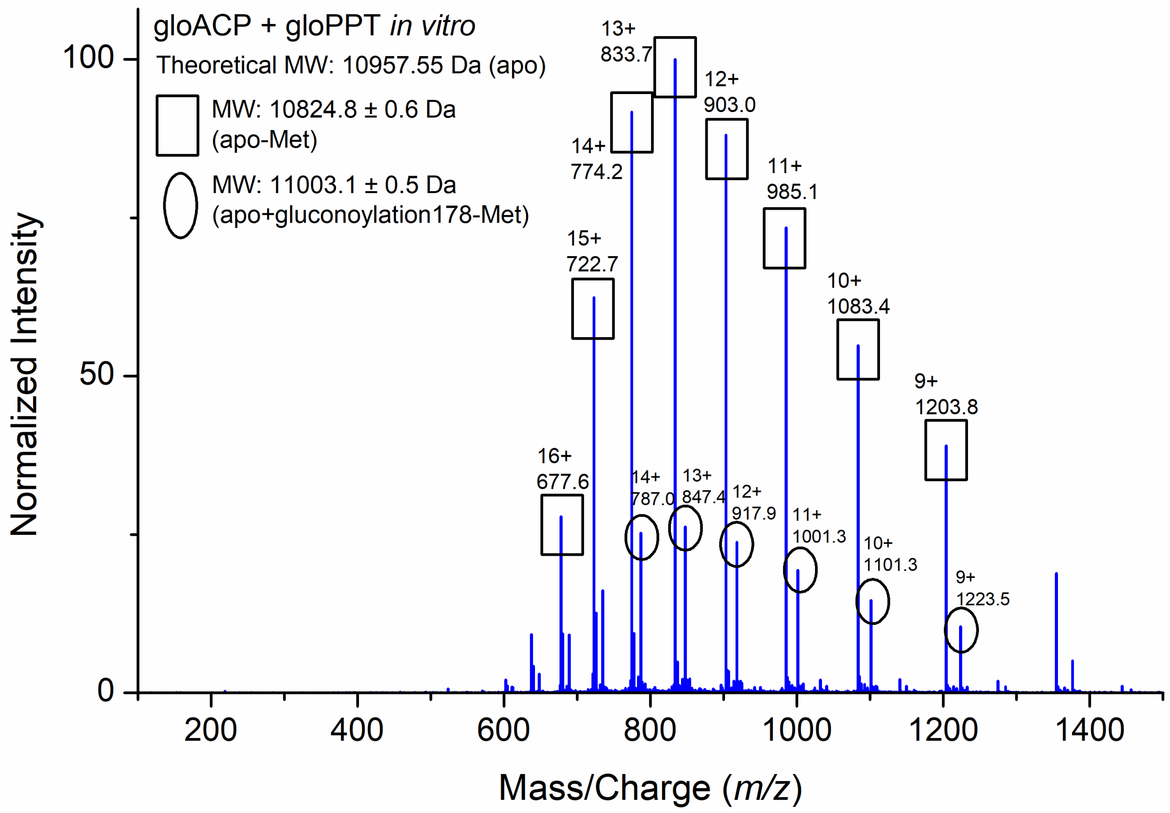


**Figure S9.** LC-MS spectrum of purified *apo*-gloACP incubated *in vitro* with separately expressed and purified gloPPT. Additional reaction components include DTT, coenzyme A, and $MgCl_2$. No phosphopantetheinylation of ACP was observed.

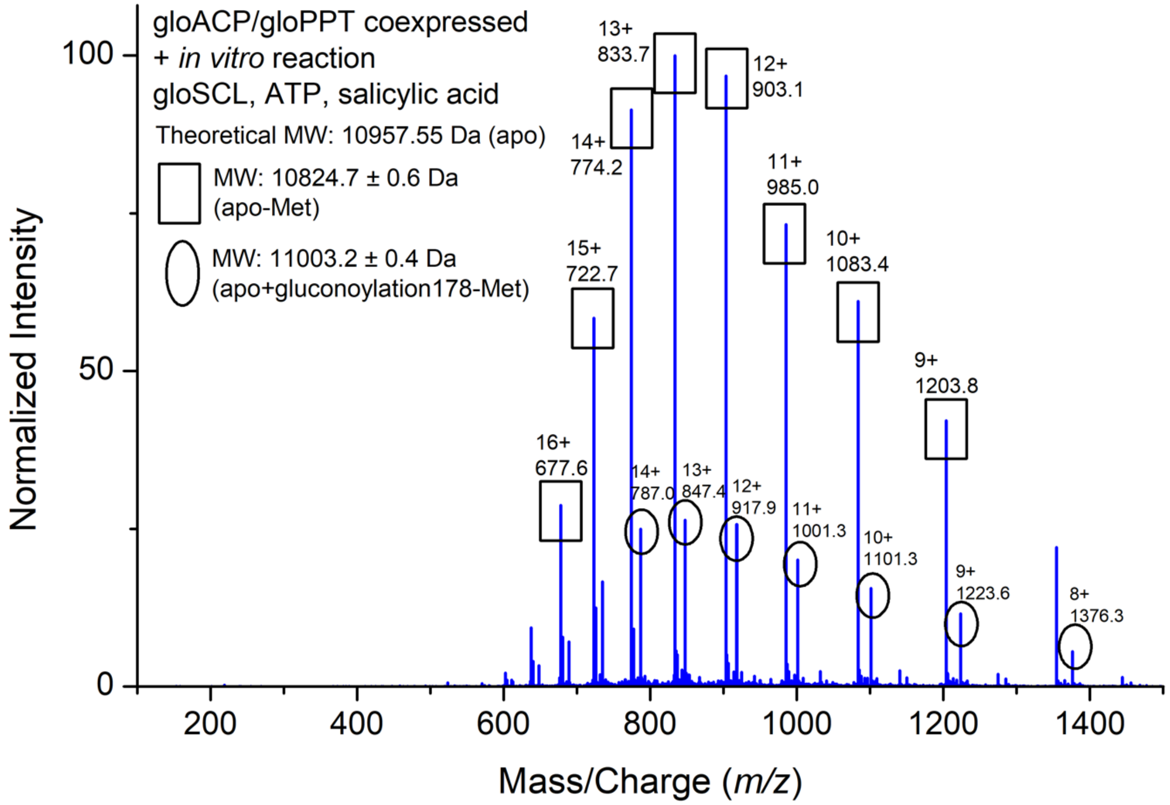


**Figure S10.** LC-MS spectrum of co-expressed/ co-purified *apo*-gloACP and gloPPT incubated with gloSCL. Additional reaction components include DTT, coenzyme A, $MgCl_2$, ATP, and salicylic acid. No phosphopantetheinylation of ACP was observed.

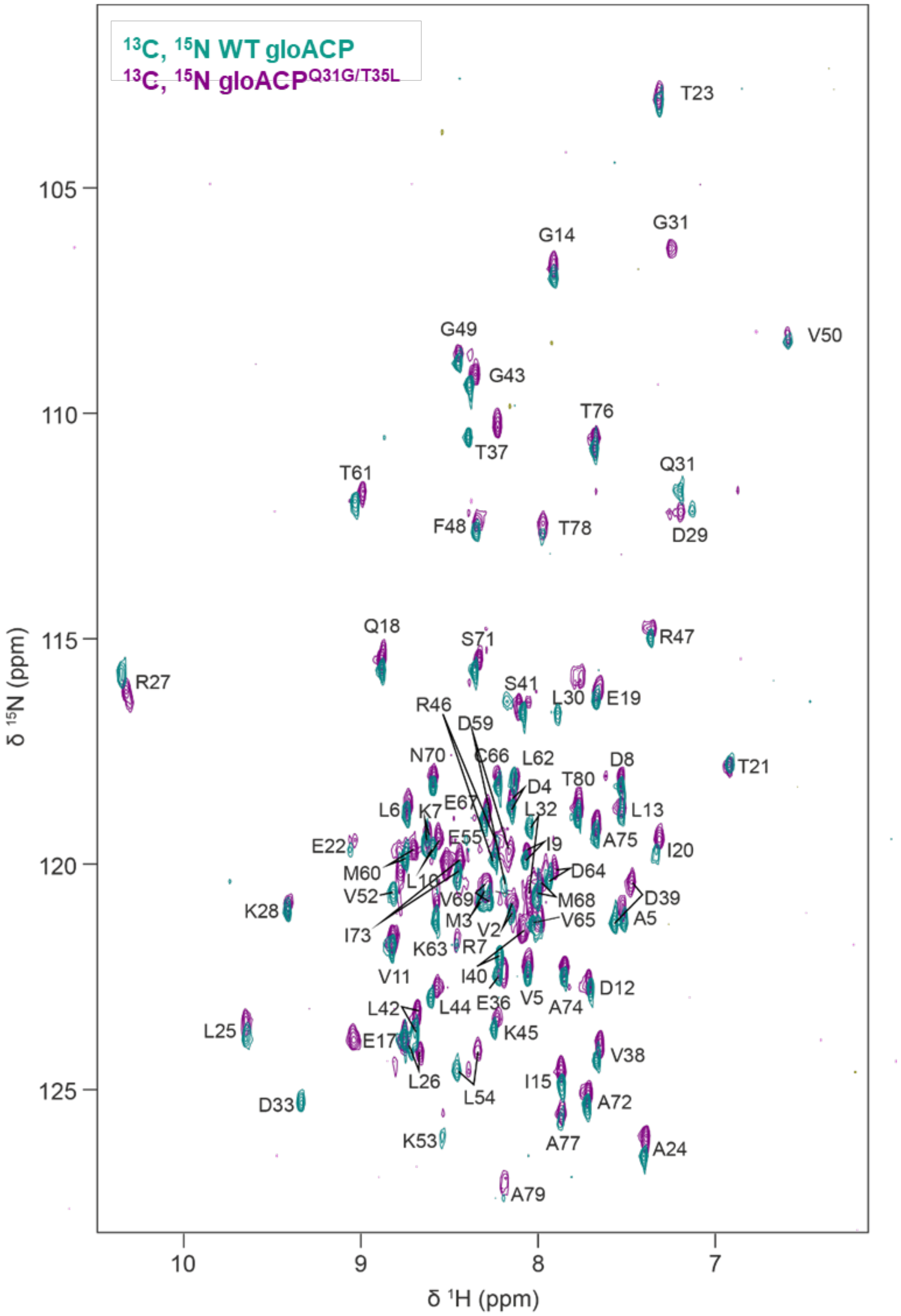


**Figure S11.** Backbone NMR assignment and overlay of the [$^{1}H$, $^{15}N$]-HSQC NMR spectra of $^{13}C$, $^{15}N$-labeled wild-type gloACP (teal) and gloACP$^{Q31G/T35L}$ mutant (purple). Assignments for backbone amide resonance are labeled in one letter amino acid code according to the *Gloeocapsa* sp. PCC 7428 Acyl Carrier Protein sequence (UniProtKB: K9XMC1).

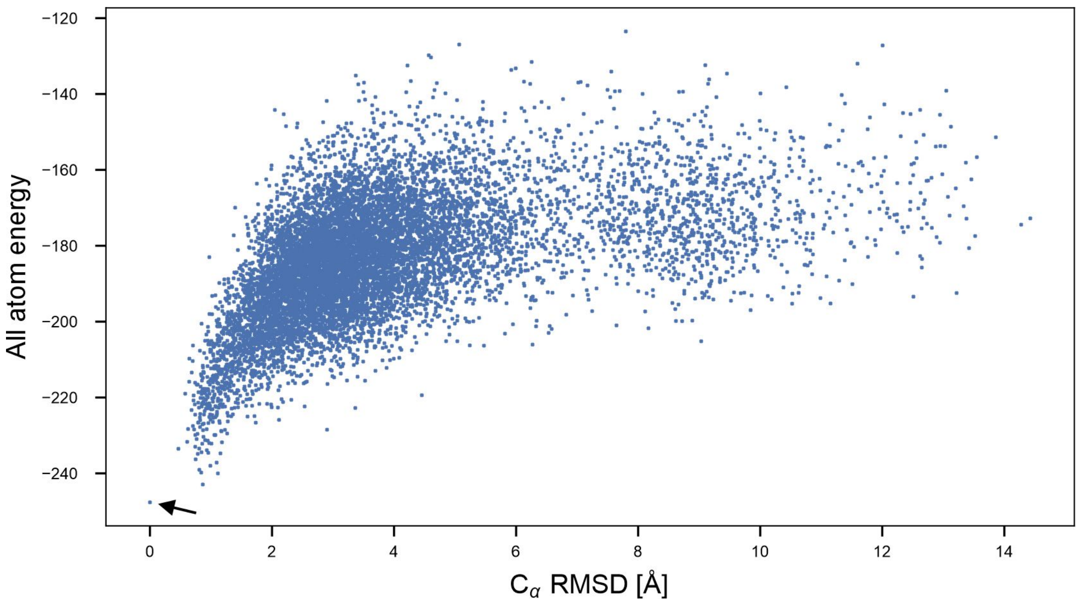


**Figure S12.** CS-Rosetta convergence plot for gloACP. Re-scored CS-Rosetta all atom energy versus Ca-RMSD relative to the model with the lowest energy (marked by an arrow) is shown. 10,000 structural models were calculated. 1292 models have a RMSD < 2 Å to the lowest energy model. Highly flexible residues at the *N*- and *C*-termini were removed for structural modelling. The ten lowest energy structural models are shown in Fig. 1A in the main paper.

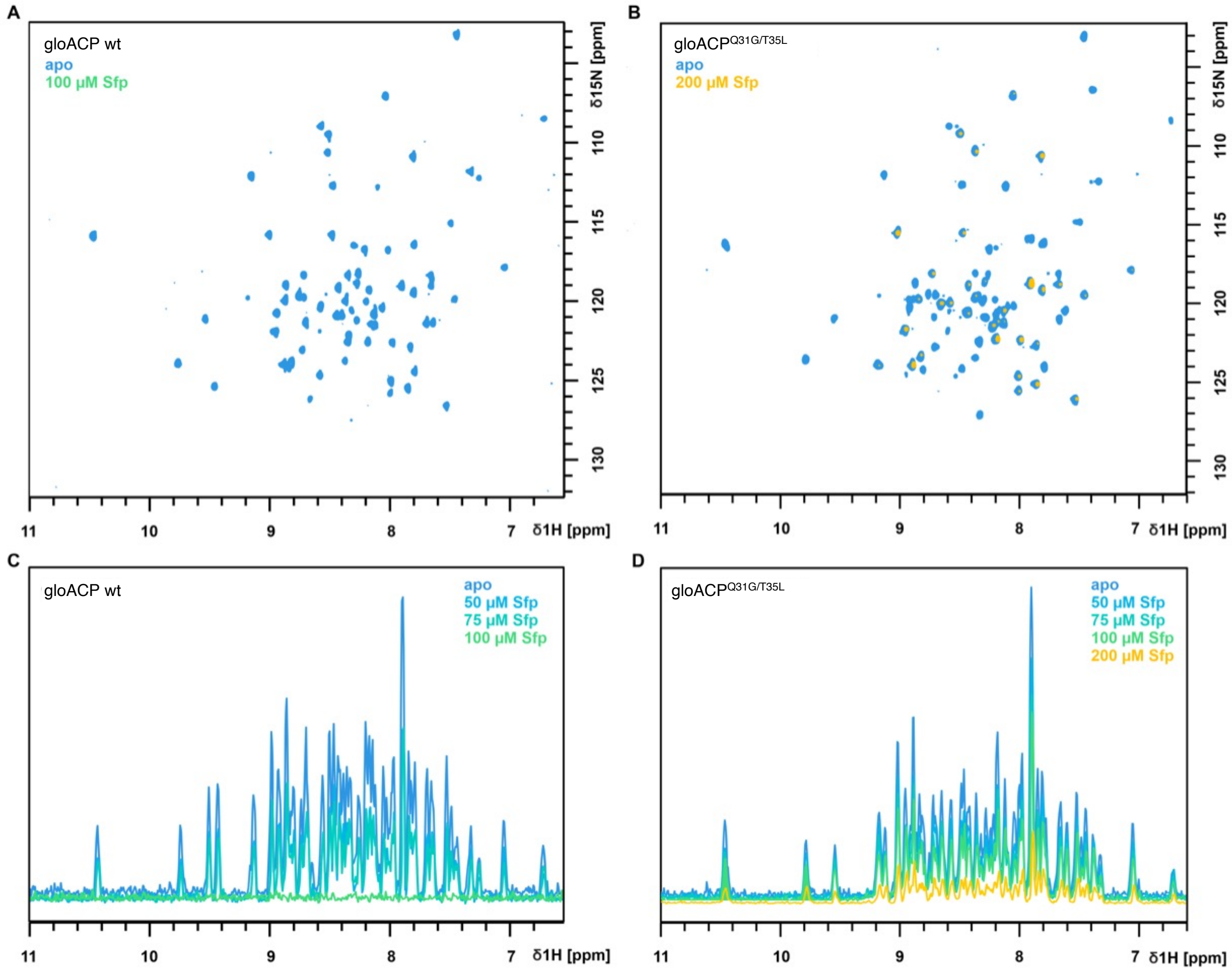


**Figure S13.** [$^1$H, $^{15}$N]-HSQC of $^{15}$N-labeled *apo*-gloACP wild-type and Q31G/T35L mutant upon titration with unlabeled Sfp.
(A) [$^1$H, $^{15}$N]-HSQC NMR spectra of $^{15}$N-labeled wild-type *apo*-gloACP in the absence (blue) and in the presence of Sfp (green).
(B) [$^1$H, $^{15}$N]-HSQC NMR spectra of $^{15}$N-labeled gloACP$^{Q31G/T35L}$ in the absence (blue) and presence of Sfp (yellow). (C, D) 1D $^1$H projections of HSQC spectra shown in (A) and (B). (C) WT *apo*-gloACP shows line broadening at significantly lower Sfp concentrations than *apo*-gloACP$^{Q31G/T35L}$ consistent with a reduced Sfp affinity in the ACP mutant.

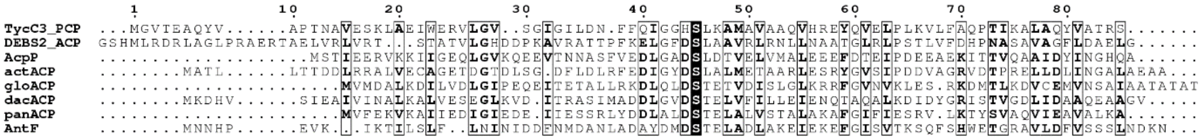


**Figure S14.** Multiple sequence alignment of selected carrier proteins known to be modified by Sfp (top four) and non-actinobacterial ACPs that demonstrate Sfp-incompatibility (bottom four, refer to Fig. 2A for more details). The serine point of attachment for the phosphopantetheine arm is highlighted in black for each sequence. Black boxes represent residues containing greater than 70% physicochemical similarity across all compared sequences.

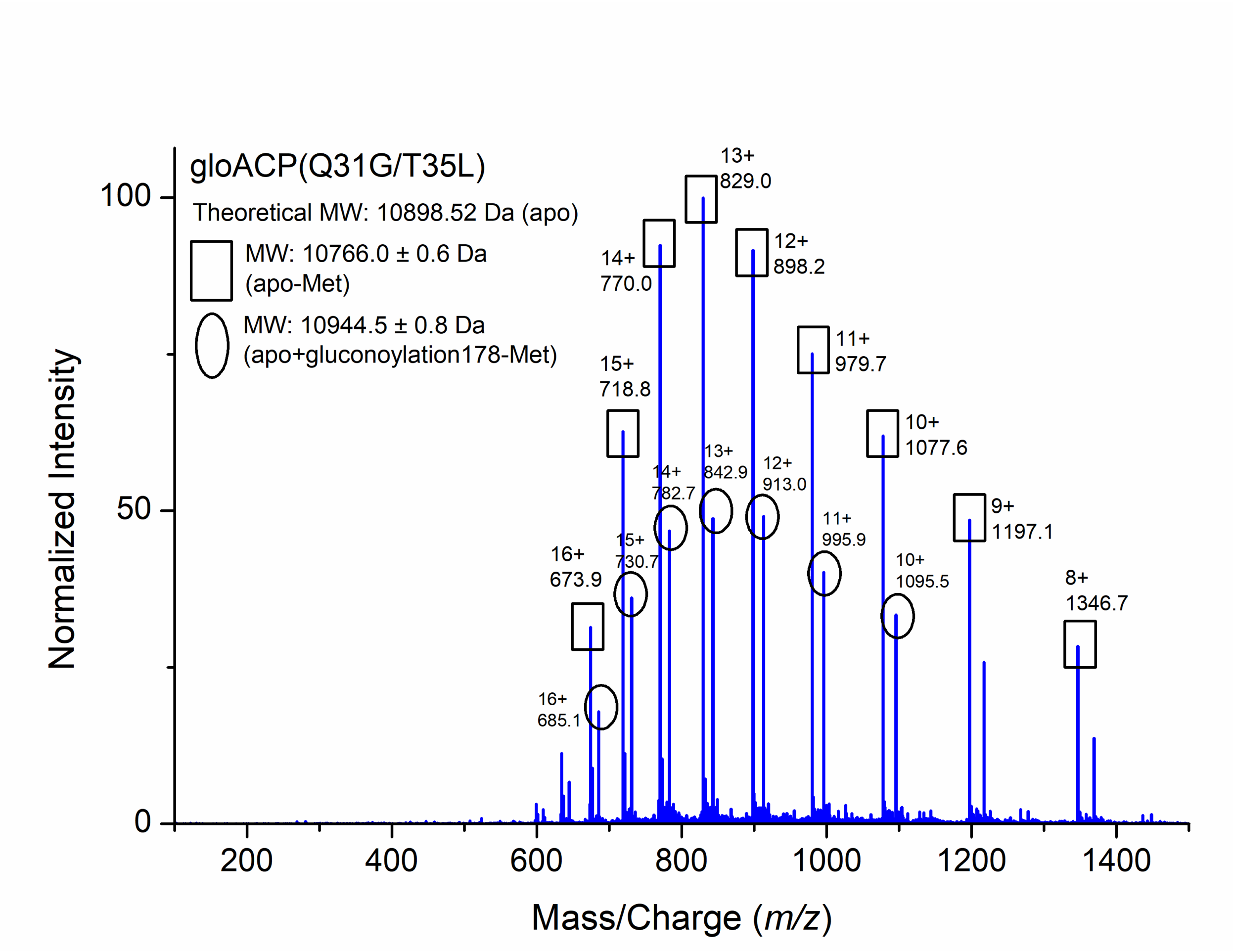


**Figure S15.** LC-MS spectrum of purified *apo*-gloACP$^{Q31G/T35L}$ heterologously expressed in *E. coli* BL21 (DE3).

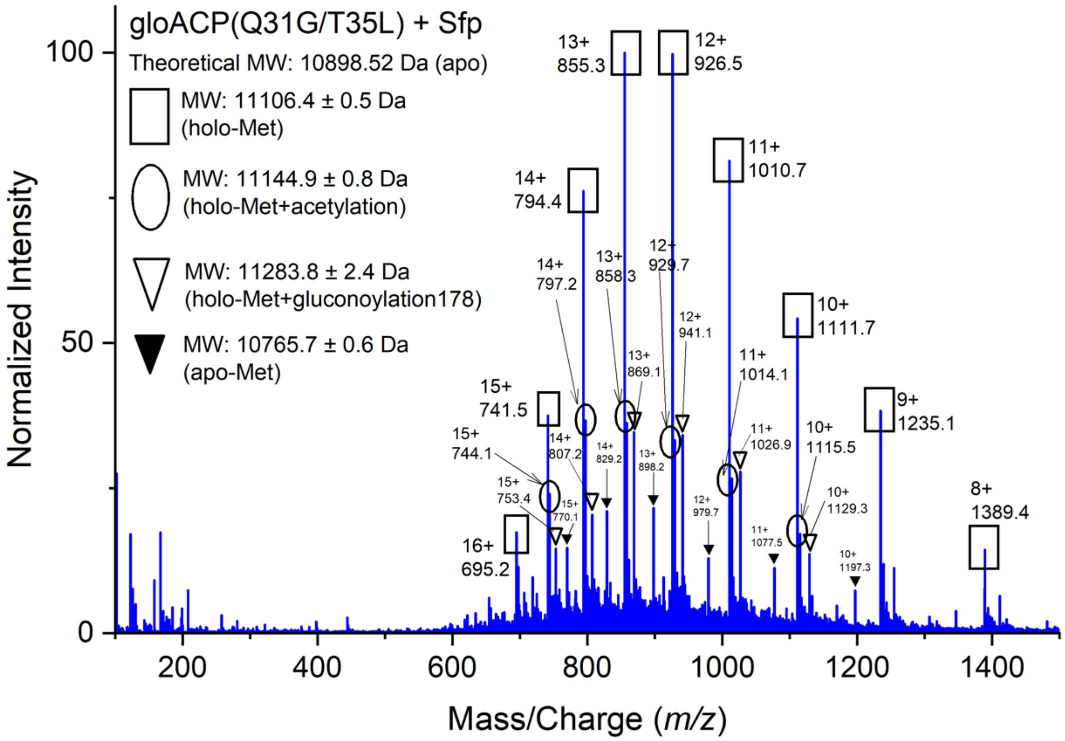


**Figure S16.** LC-MS spectrum of gloACP$^{Q31G/T35L}$ reacted with Sfp via heterologous expression in *E. coli* BAP1 *without* additional *in vitro* incubation with Sfp (and DTT, coenzyme A, and $MgCl_2$), showing successful conversion to *holo*-gloACP$^{Q31G/T35L}$.

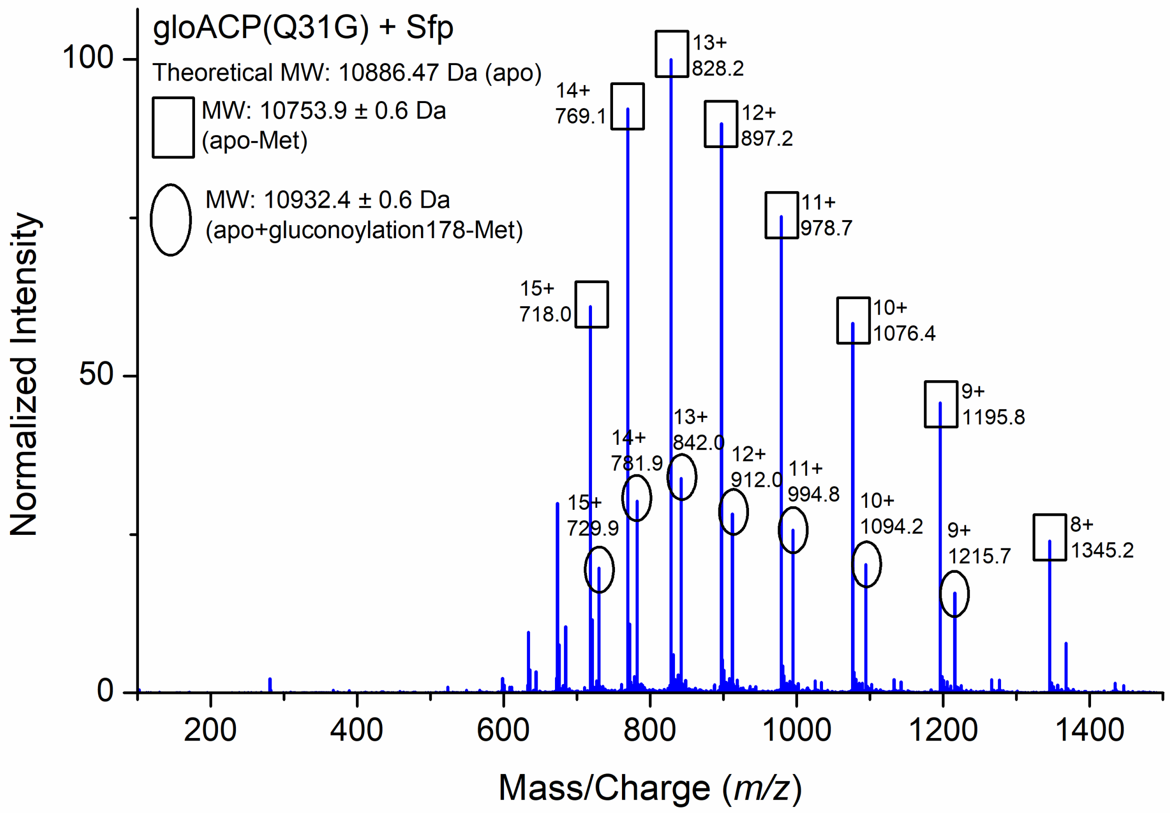


**Figure S17.** LC-MS spectrum of *apo*-gloACP$^{Q31G}$. The protein was expressed and purified from *E. coli* BAP1 and additionally incubated *in vitro* with Sfp, DTT, coenzyme A, and $MgCl_2$. No phosphopantetheinylation of ACP was observed.

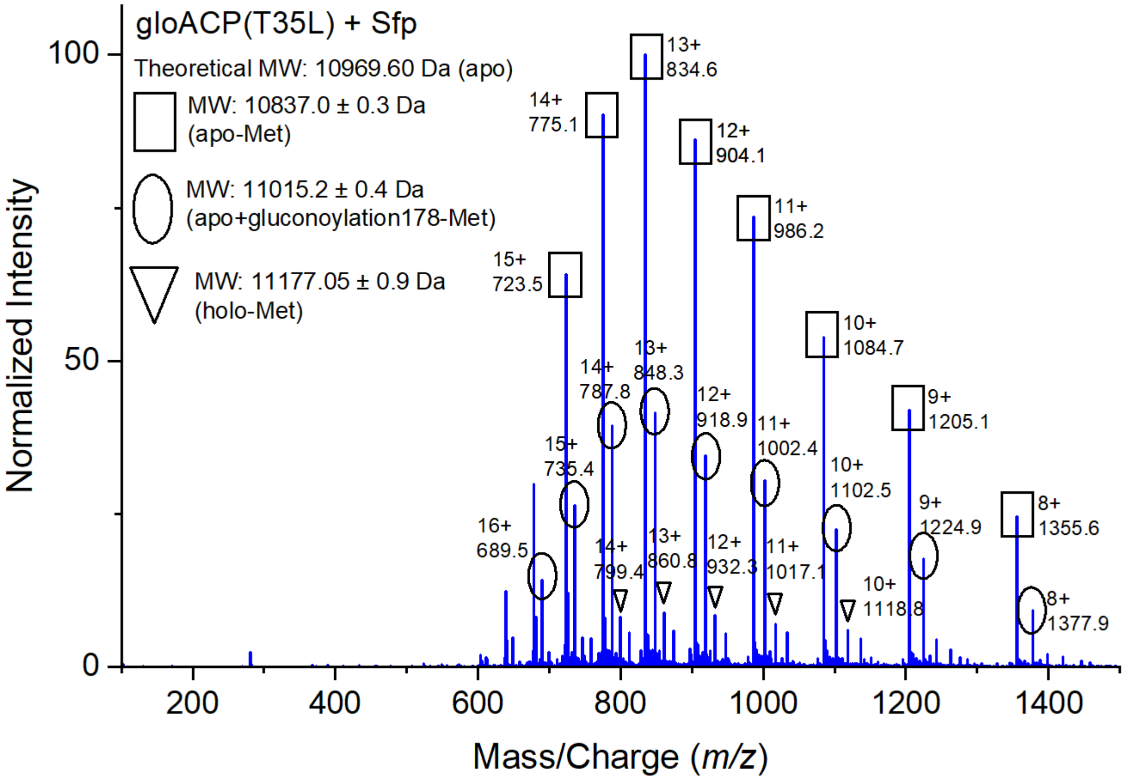


**Figure S18.** LC-MS spectrum of gloACP$^{T35L}$ incubated with Sfp, resulting in minimal *holo*-gloACP$^{T35L}$ formation. GloACP$^{T35L}$ was expressed and purified from *E. coli* BAP1 and additionally incubated *in vitro* with Sfp, DTT, coenzyme A, and $MgCl_2$.

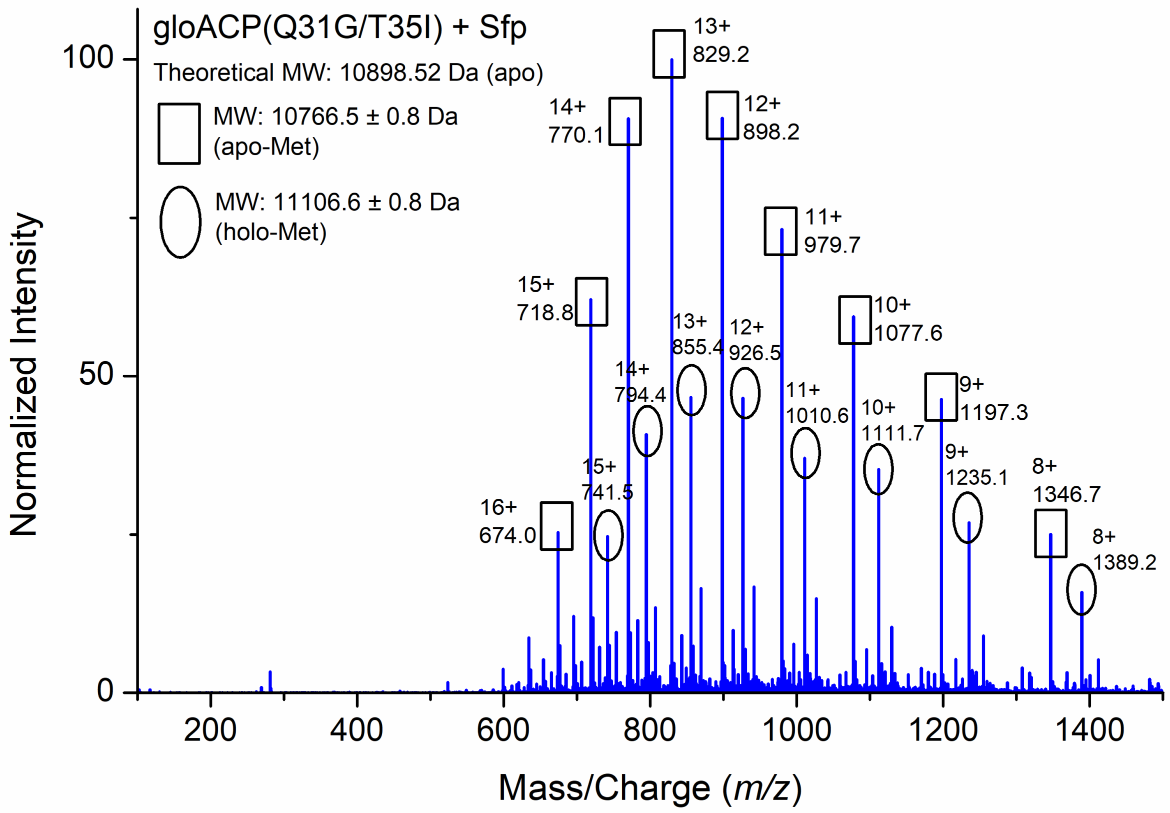


**Figure S19.** LC-MS spectrum of *apo*-gloACP$^{Q31G/T35I}$ incubated with Sfp results in *holo*-gloACP$^{Q31G/T35I}$ formation. GloACP$^{Q31G/T35I}$ was first expressed and purified from *E. coli* BAP1 and additionally incubated *in vitro* with Sfp, DTT, coenzyme A, and $MgCl_2$.

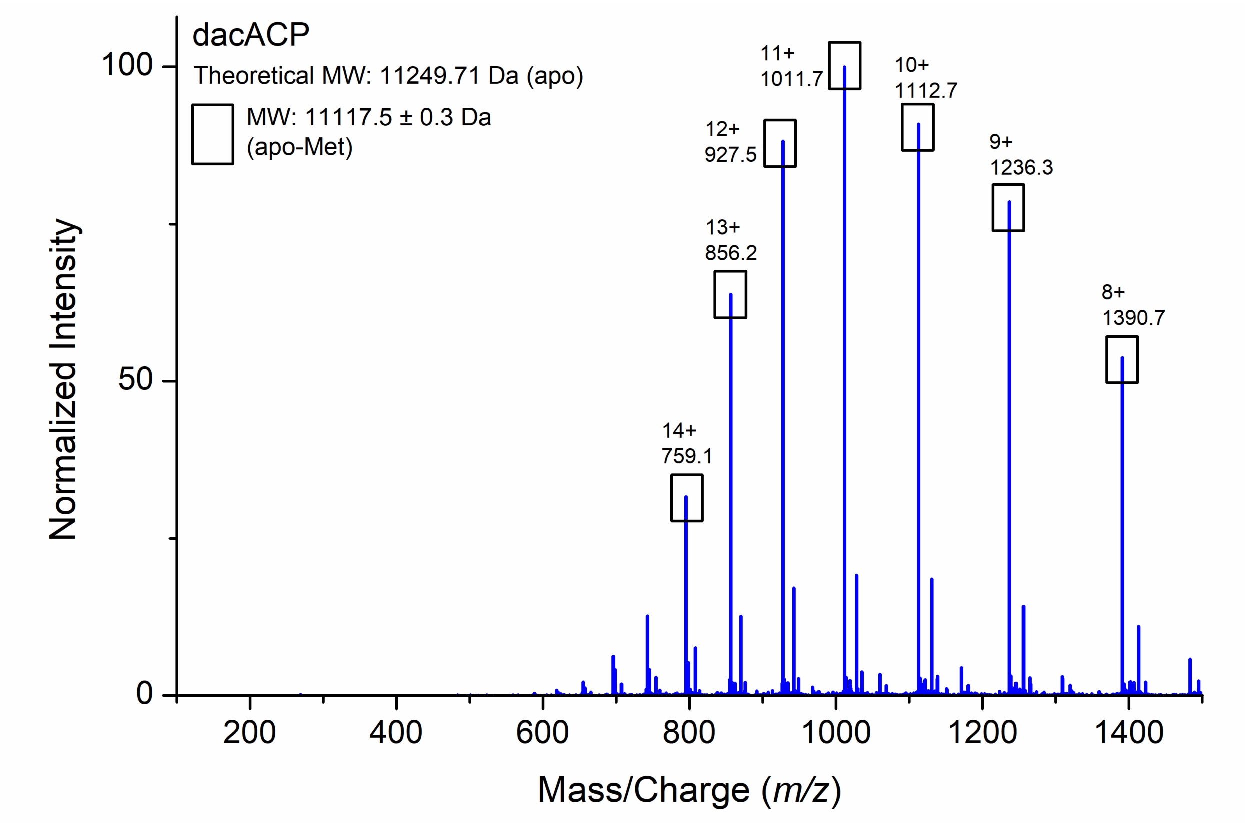


**Figure S20.** LC-MS spectrum of purified *apo*-dacACP.

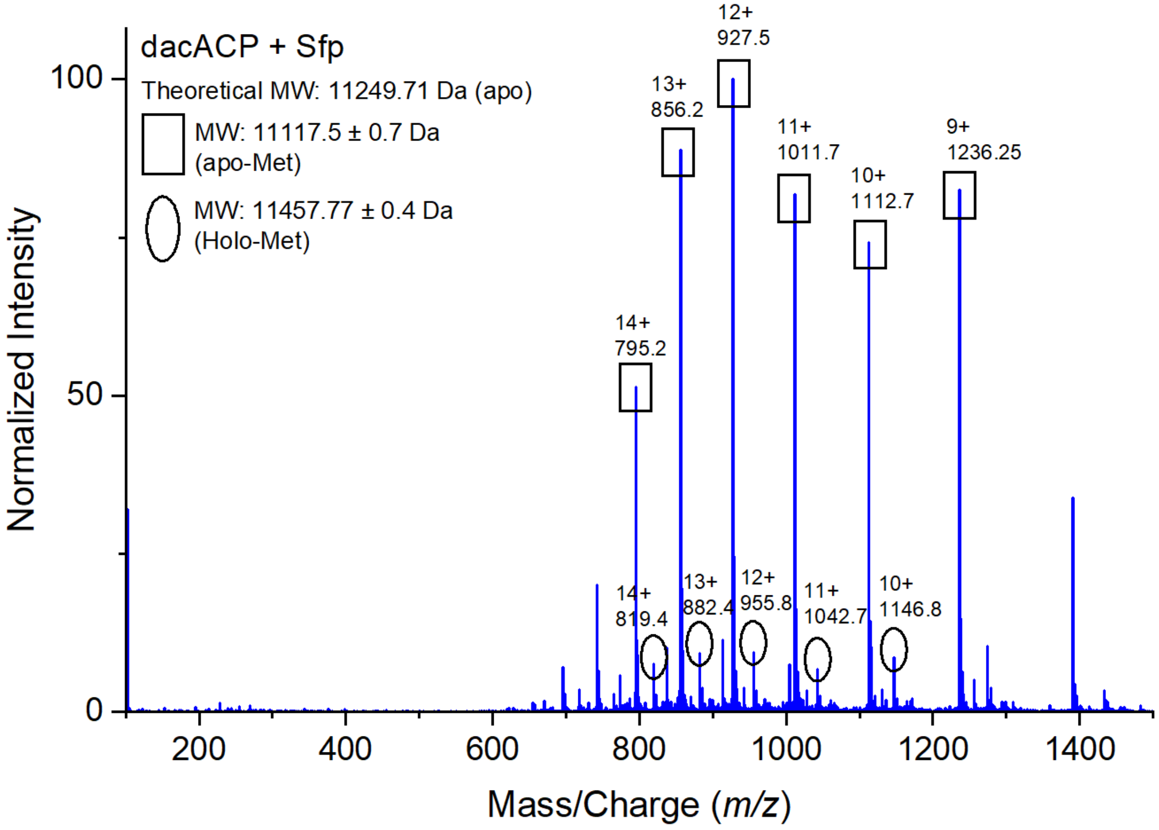


**Figure S21.** LC-MS spectrum of purified dacACP incubated with Sfp resulting in minimal *holo*-dacACP formation. DacACP was first expressed and purified from *E. coli* BAP1 and additionally incubated *in vitro* with Sfp, DTT, coenzyme A, and $MgCl_2$.

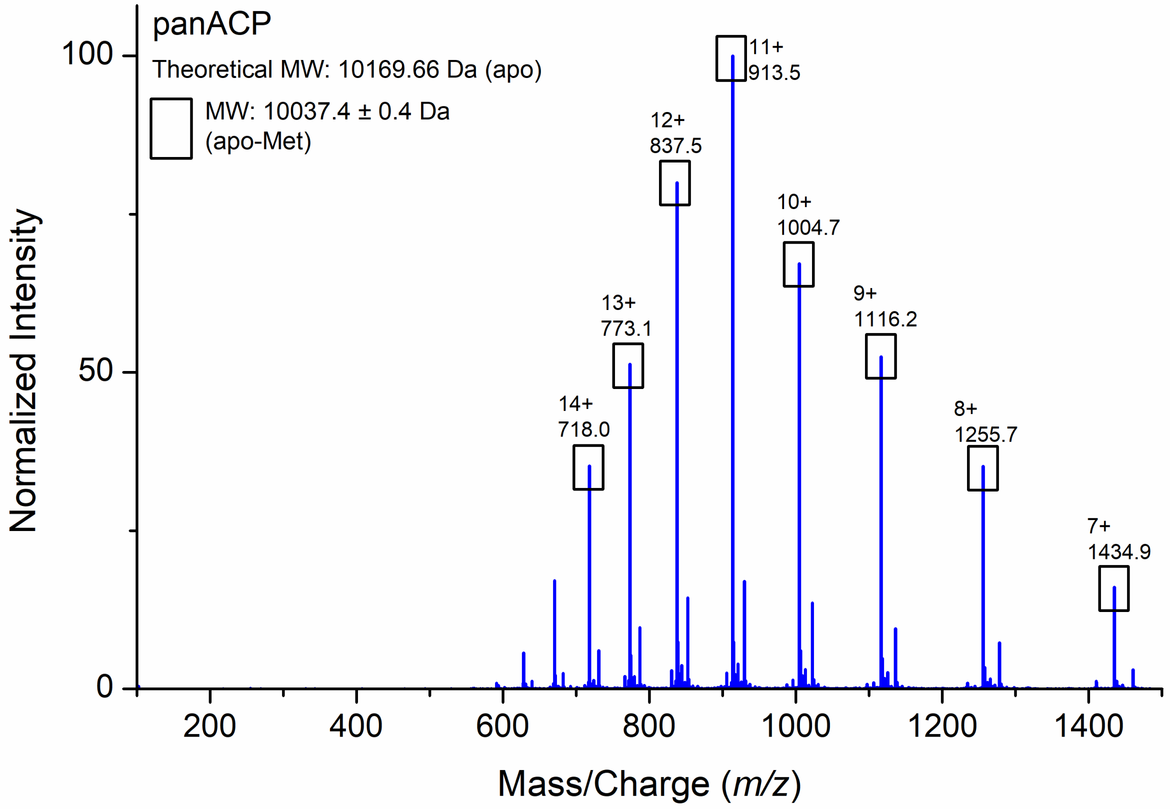


**Figure S22.** LC-MS spectrum of purified *apo*-panACP.

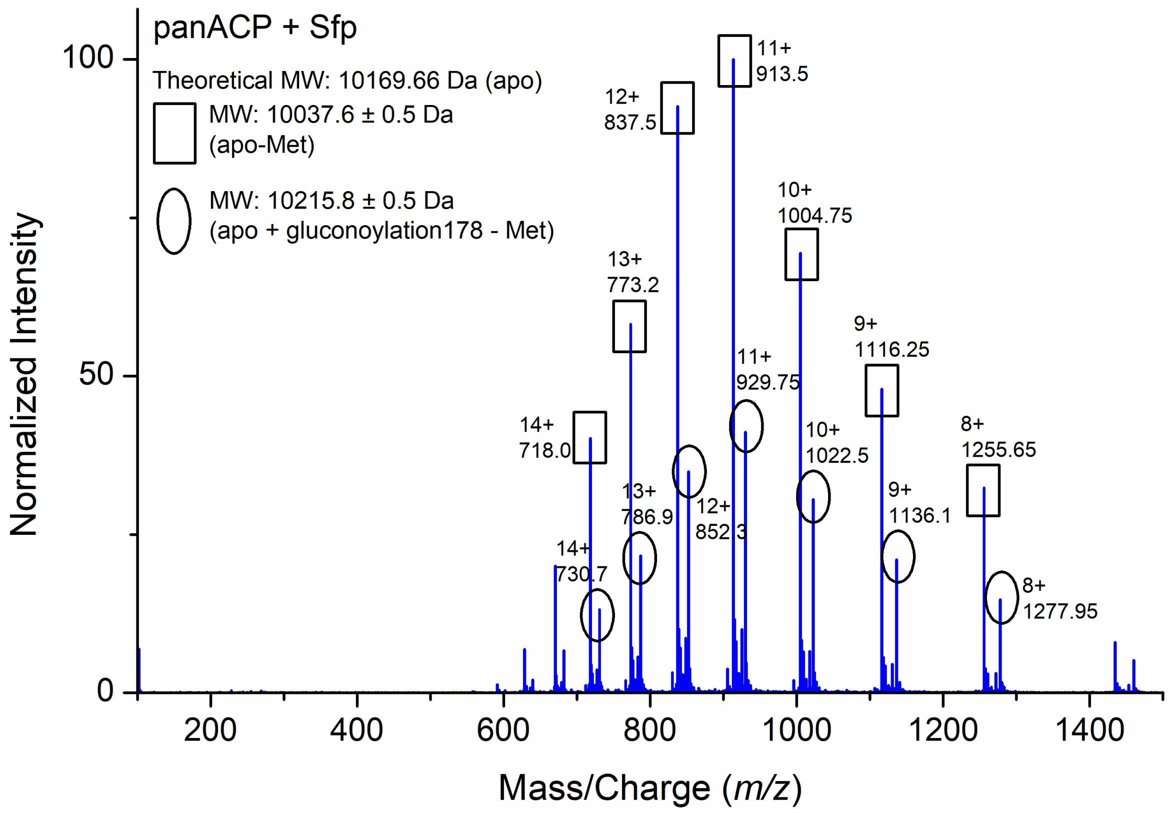


**Figure S23.** LC-MS spectrum of purified panACP incubated with Sfp. PanACP was expressed and purified from *E. coli* BAP1 and subsequently additionally incubated with Sfp, DTT, coenzyme A, and $MgCl_2$. No phosphopantetheinylation of ACP was observed.

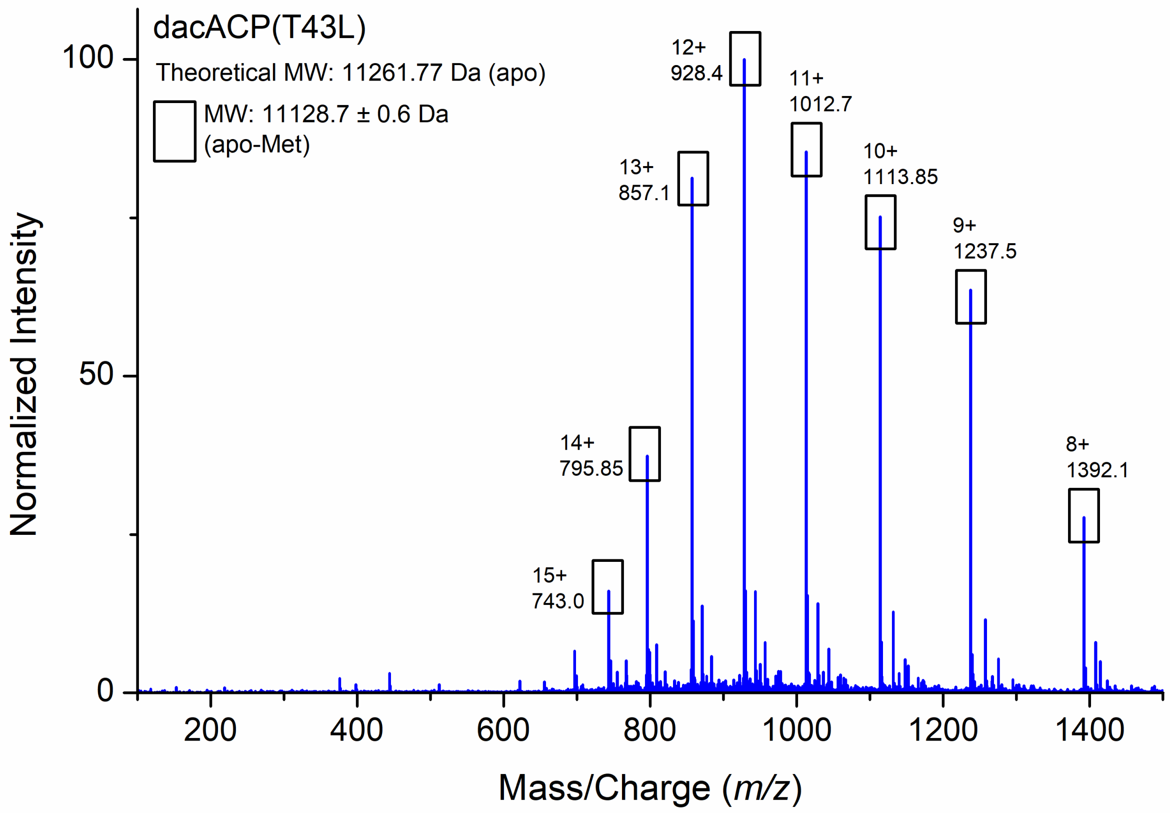


**Figure S24.** LC-MS spectrum of purified *apo*-dacACP$^{T43L}$.

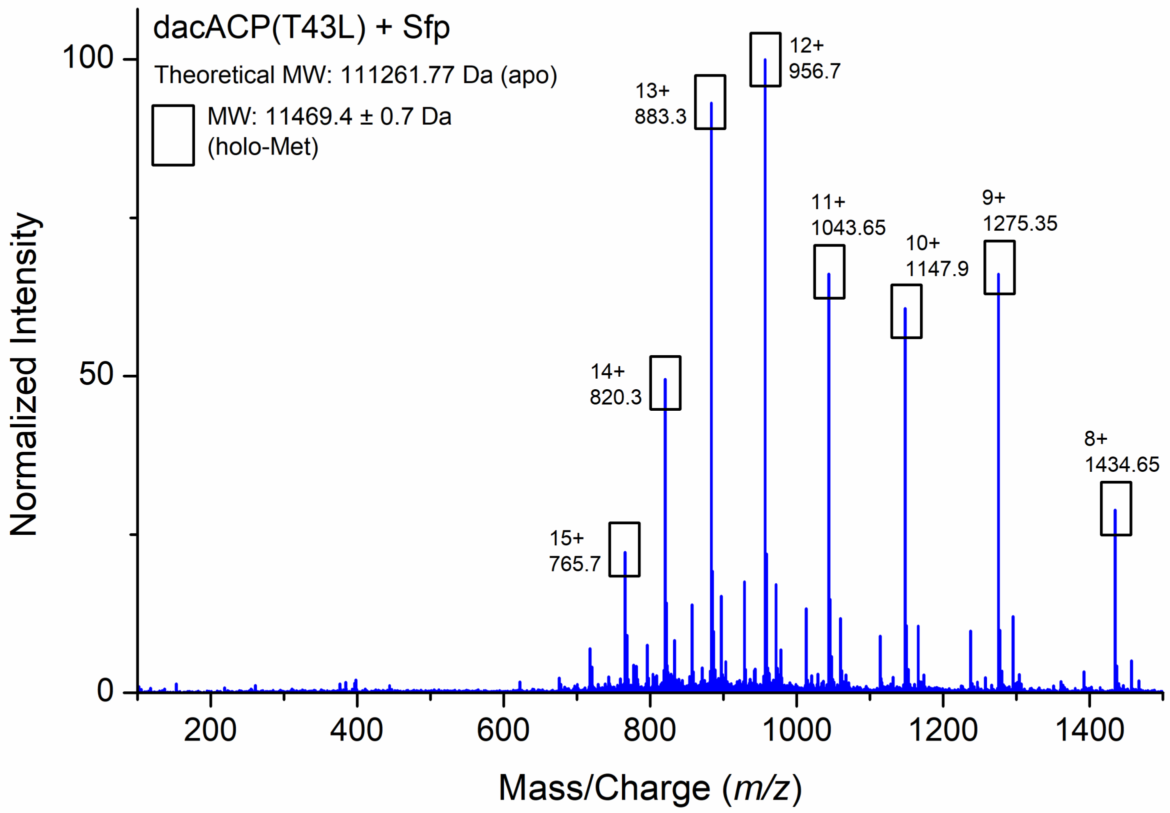


**Figure S25.** LC-MS spectrum of purified *holo*-dacACP$^{T43L}$ upon incubation with Sfp. DacACP$^{T43L}$ was expressed and purified from *E. coli* BAP1 and additionally incubated *in vitro* with Sfp, DTT, coenzyme A, and $MgCl_2$. Successful conversion to *holo*-dacACP$^{T43L}$ was observed.

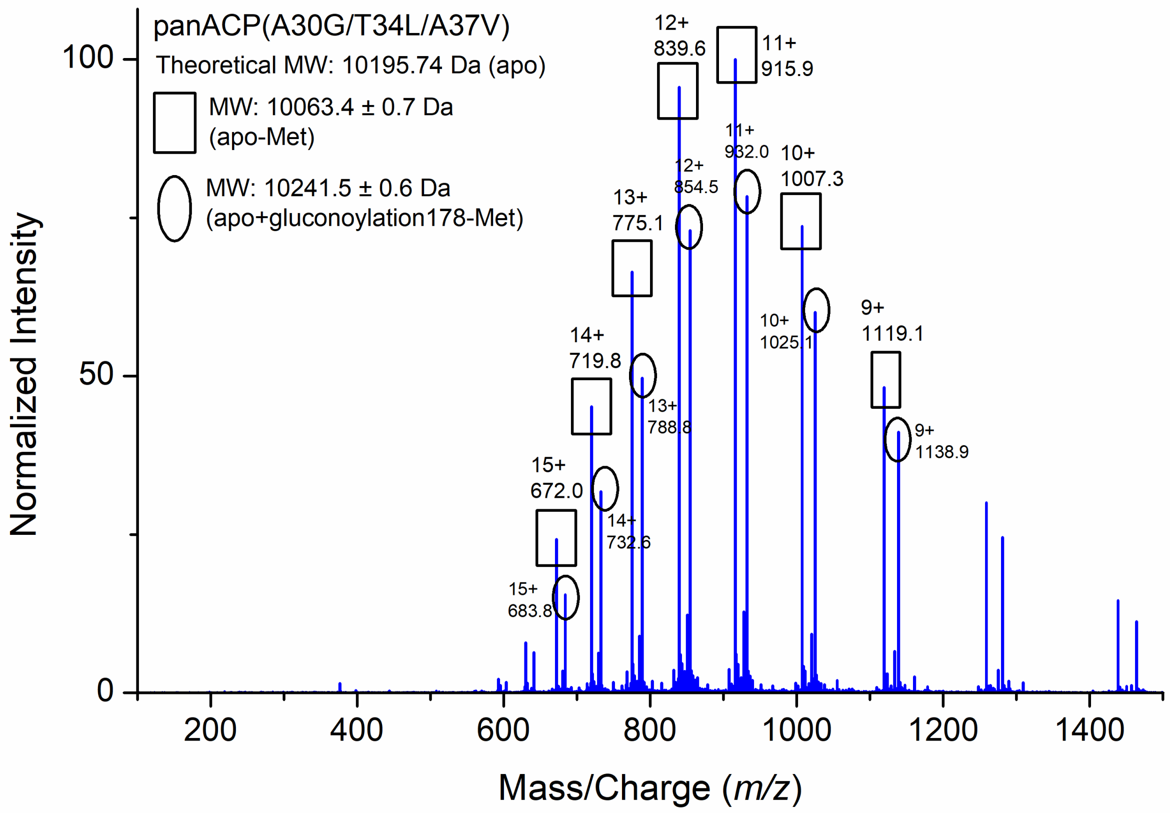


**Figure S26.** LC-MS spectrum of purified *apo*-panACP$^{A30G/T34L/A37V}$.

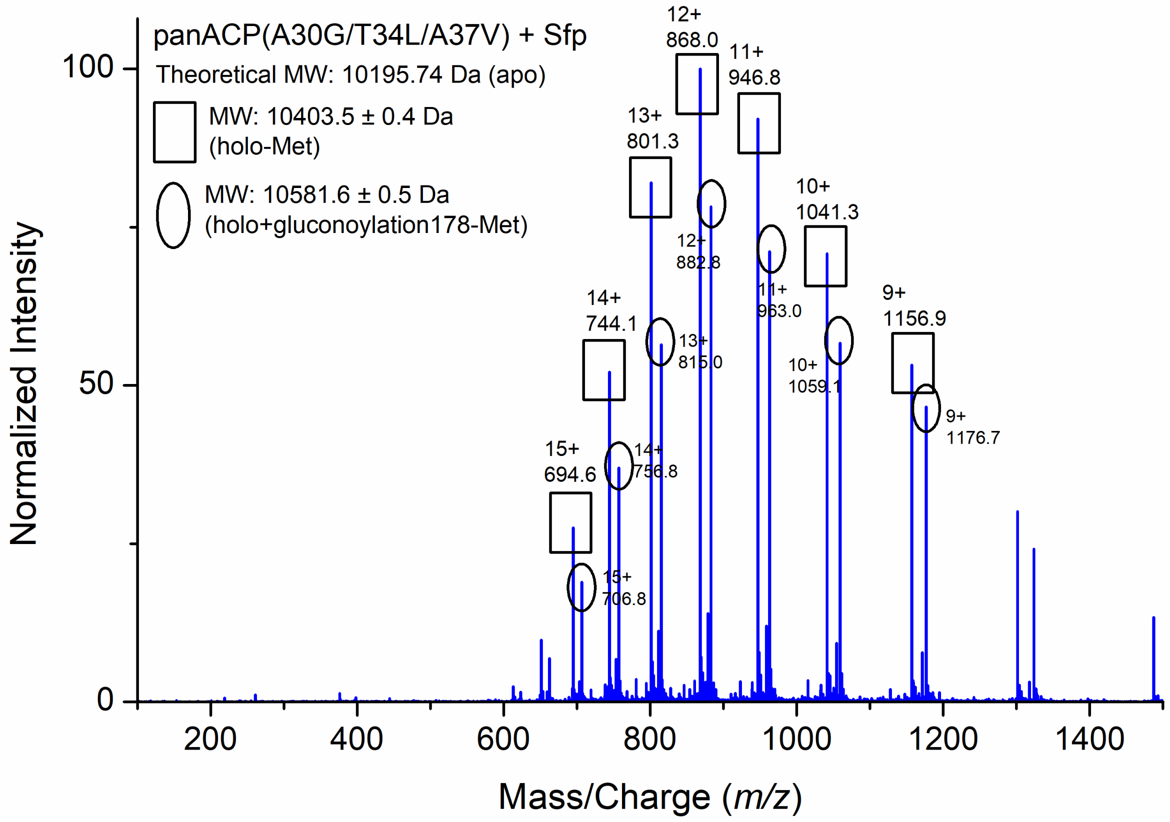


**Figure S27.** LC-MS spectrum of purified *holo*-panACP$^{A30G/T34L/A37V}$ upon incubation with Sfp. PanACP$^{A30G/T34L/A37V}$ was first expressed and purified from *E. coli* BAP1 and additionally incubated *in vitro* with Sfp, DTT, coenzyme A, and $MgCl_2$. Successful conversion to *holo*-panACP$^{A30G/T34L/A37V}$ was observed.

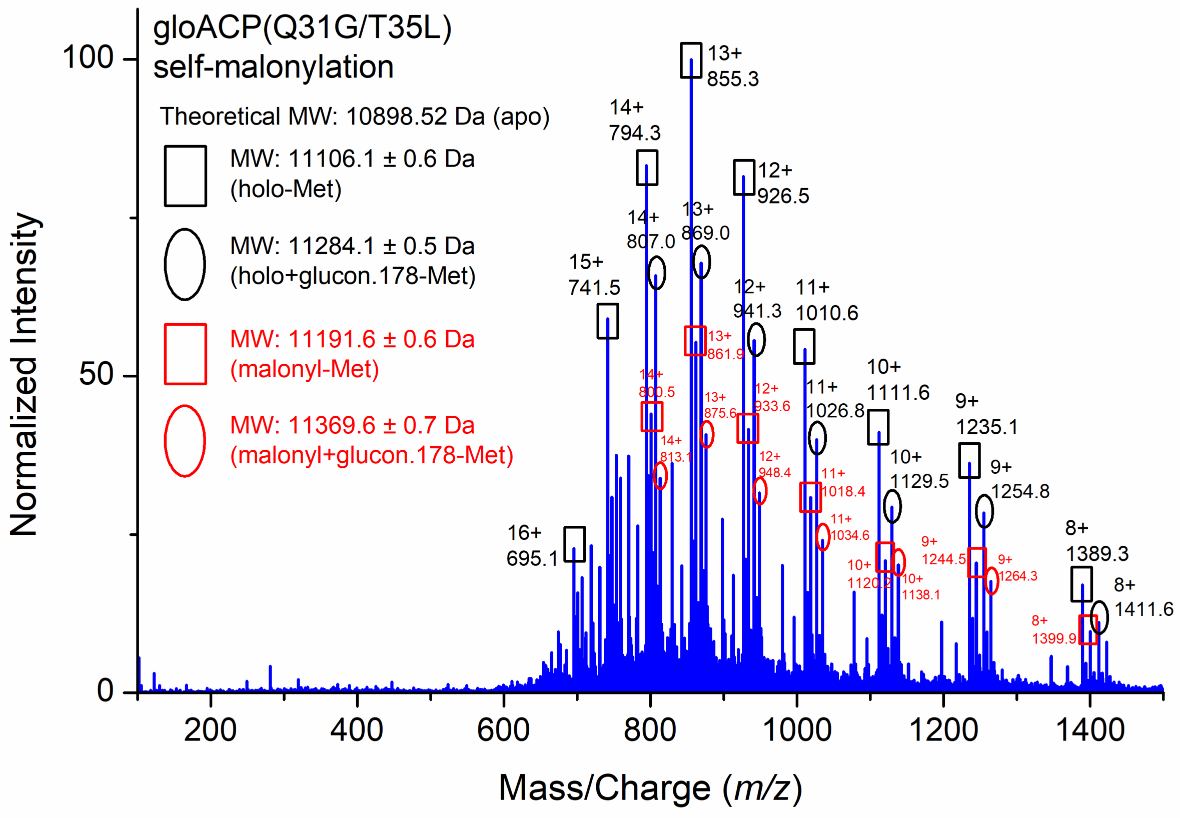


**Figure S28.** LC-MS spectrum of malonyl-gloACP$^{Q31G/T35L}$ from self-malonylation of *holo*-gloACP$^{Q31G/T35L}$ upon incubation with malonyl CoA.

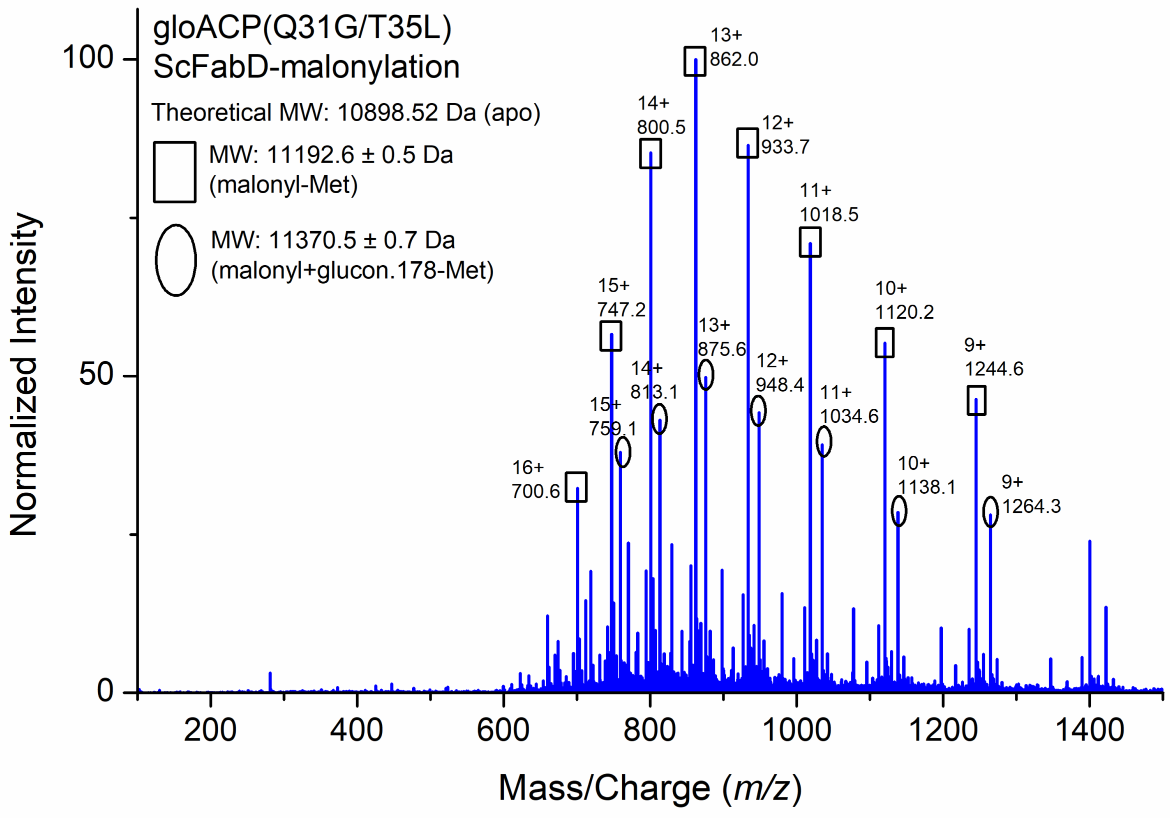


**Figure S29.** LC-MS spectrum of malonyl-gloACP$^{Q31G/T35L}$ after ScFabD-catalyzed malonlylation of *holo*-gloACP$^{Q31G/T35L}$.

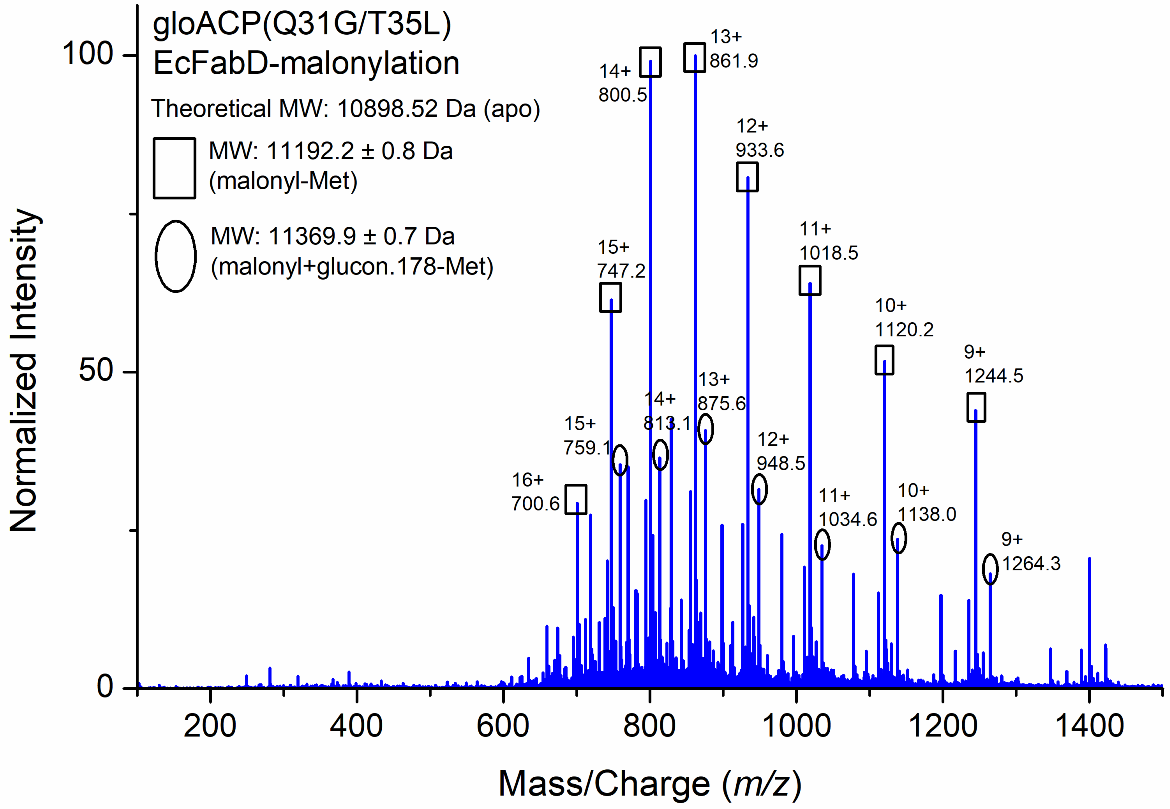


**Figure S30.** LC-MS spectrum of malonyl-gloACP$^{Q31G/T35L}$ after EcFabD-catalyzed malonlylation of *holo*-gloACP$^{Q31G/T35L}$.

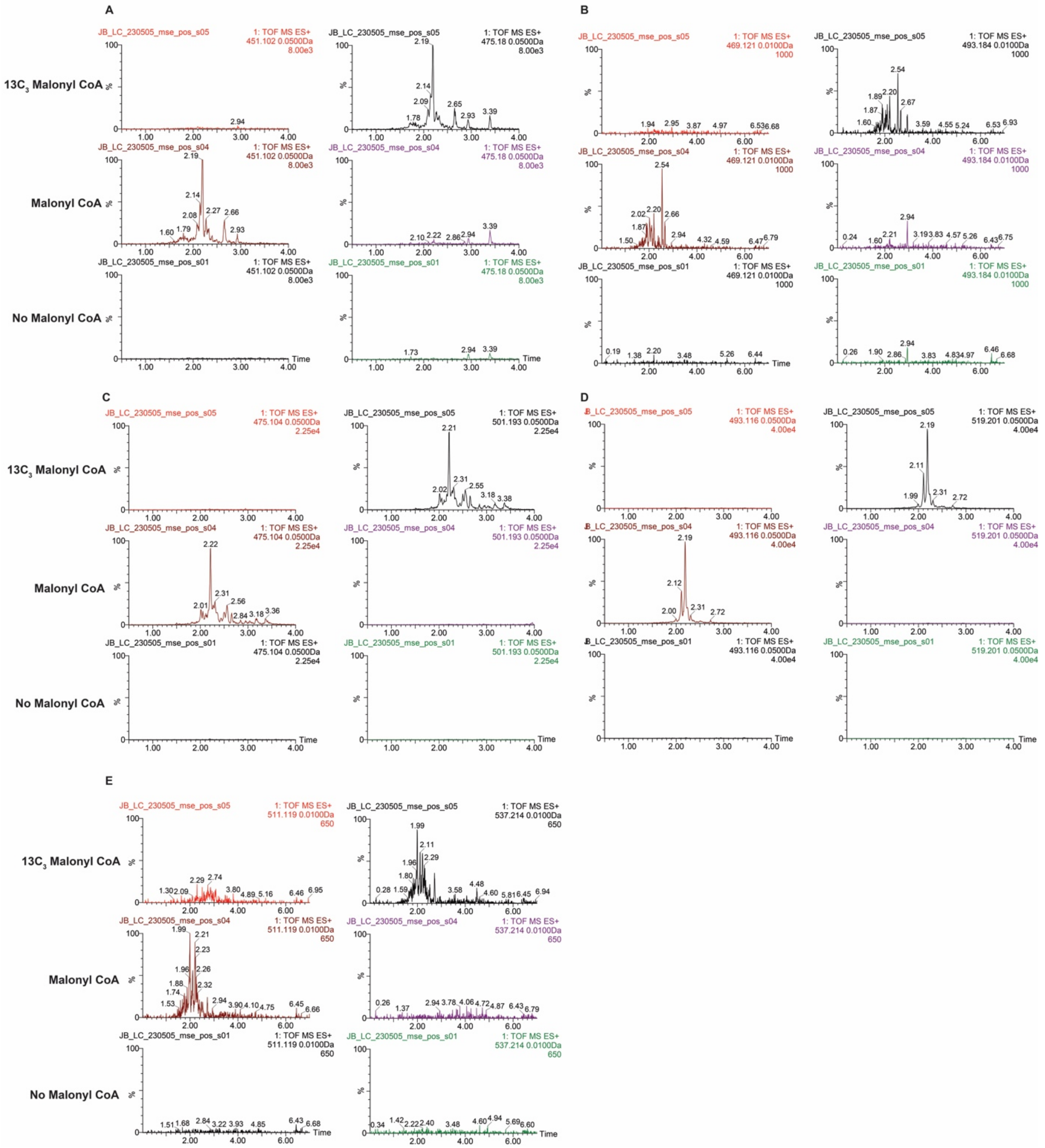


**Figure S31.** Full extracted high resolution ion chromatograms of mass peaks of *m/z* 451, 469, 475, 493, and 511 produced by the reconstituted core gloPKS in the absence of gloSCL/ salicylate (compare Fig. 3). (A) Malonyl-CoA derived mass peak of *m/z* 451, which shifts 24 amu to 475 upon isotopic labeling with $^{13}C$-malonyl-CoA. (B) Malonyl-CoA derived mass peak of *m/z* 469, which shifts 24 amu to 493 upon isotopic labeling. (C) Malonyl-CoA derived mass peak of *m/z* 475, which shifts 26 amu to 501 upon isotopic labeling. (D) Malonyl-CoA derived mass peak of *m/z* 493, which shifts 26 amu to 519 upon isotopic labeling. (E) Malonyl-CoA derived mass peak of *m/z* 511, which shifts 26 amu to 537 upon isotopic labeling.

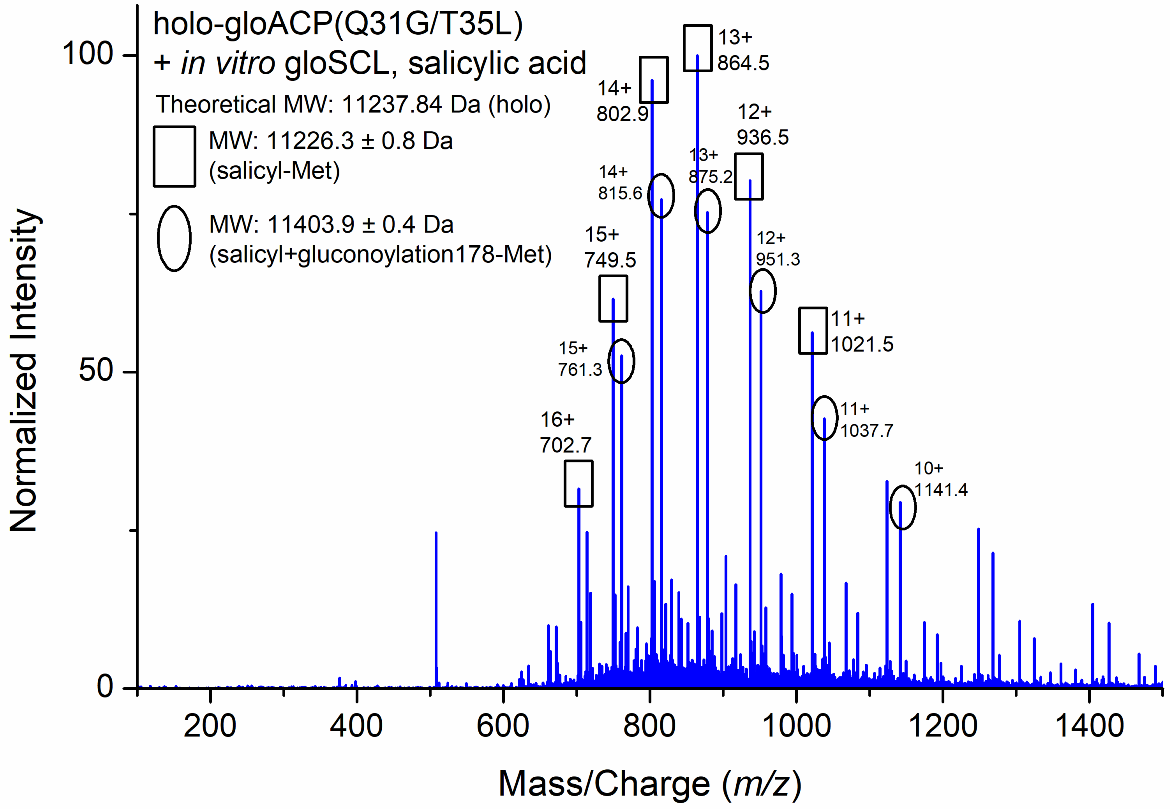


**Figure S32.** LC-MS spectrum of purified salicyl-gloACP$^{Q31G/T35L}$ produced upon incubation of *holo*-gloACP$^{Q31G/T35L}$ with gloSCL, salicylic acid, ATP, and tris(2-carboxyethyl)phosphine (TCEP).

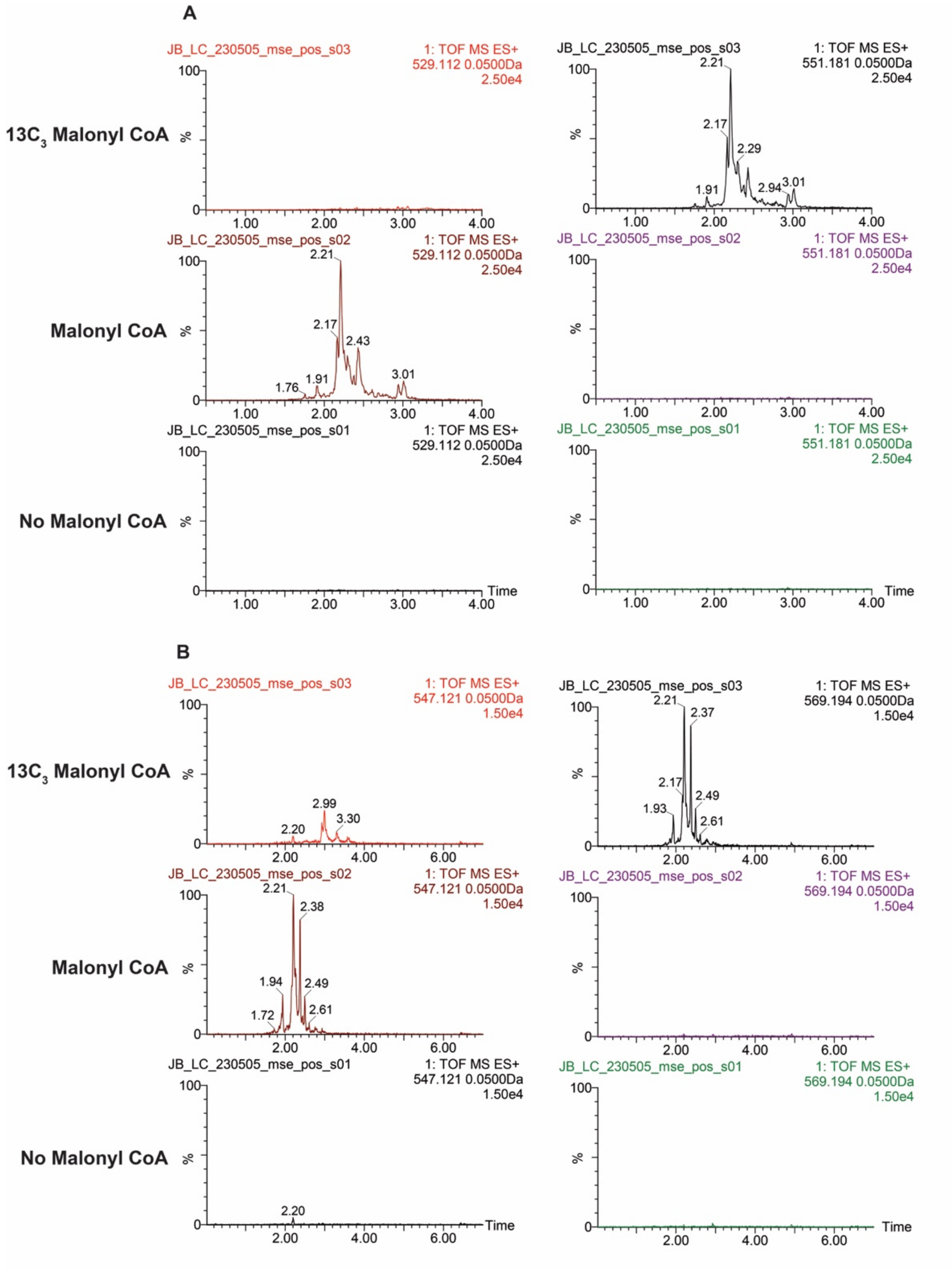


**Figure S33**. Full extracted high resolution ion chromatograms of product peaks of *m/z* 529 and 547 produced by the reconstituted core gloPKS in the presence of gloSCL/ salicylate (compare Fig. 4). (A) Salicyl-incorporated polyketides of *m/z* 529, which shift 22 amu to 551 upon isotopic labeling. (B) Salicyl-incorporated polyketides of *m/z* 547, which shift 22 amu to 569 upon isotopic labeling.
